# Estrogen controls mTOR signaling and mitochondrial function via WNT4 in lobular carcinoma cells

**DOI:** 10.1101/684530

**Authors:** Madeleine T. Shackleford, Deviyani M. Rao, Evelyn K. Bordeaux, Hannah M. Hicks, Steffi Oesterreich, Matthew J. Sikora

## Abstract

Invasive lobular carcinoma of the breast (ILC) is strongly estrogen-driven and represents a unique context for estrogen receptor (ER) signaling. In ILC, ER controls the expression of the Wnt ligand WNT4, which is critical for endocrine response and anti-estrogen resistance. However, signaling mediated by WNT4 is poorly understood. We utilized reverse phase protein array (RPPA) to characterize ER and WNT4-driven signaling in ILC cells and identified that WNT4 mediates downstream mTOR signaling via S6K. Additionally, independent of mTOR/S6K, ER and WNT4 control levels of MCL-1, which alters mitochondrial function. In this context, WNT4 knockdown caused decreased ATP production and increased mitochondrial fragmentation. WNT4 regulation of both mTOR signaling and MCL-1 were also observed in antiestrogen resistant models of ILC. Further, we identified that high *WNT4* expression is associated with similar mTOR pathway activation in serous ovarian cancer tumors, suggesting that WNT4 signaling is important in multiple tumor types. The identified downstream pathways offer insight in to WNT4 signaling and represent potential targets to overcome anti-estrogen resistance for patients with ILC.

## INTRODUCTION

Invasive lobular carcinoma of the breast (ILC) is the second most common histologic subtype of breast cancer [2–4]. Classically, ILC is characterized by small, linear chain-forming neoplastic cells which invade the mammary ducts, stroma, and adipose tissue in a characteristic single-file pattern. As such, ILC is often difficult to detect via physical exam and mammography, resulting in a later and more advanced diagnosis. This is also associated with unique sites of metastasis [5, 6] and more challenging surgical management [7, 8]. Biomarkers for ILC are consistent with the hormone-dependent ‘Luminal A’ molecular subtype (eg. estrogen receptor (ER) and progesterone receptor positive, human epidermal growth factor receptor 2 (HER2) negative) [2, 4]. About 95% of ILC tumors express ER [4]; ILC also appear to be exquisitely sensitive to and dependent on the steroid hormone estrogen. For example, estrogen-based hormone replacement therapy (with or without progestin) more strongly increases the incidence of ILC versus the more common invasive ductal carcinoma (IDC) [3].Taken together, these observations support the current paradigm that patients with ILC are ideal candidates for treatment with anti-estrogen therapies. However, retrospective studies suggest patients with ILC may not receive similar benefit from anti-estrogens as patients with IDC, ie. poorer outcomes with adjuvant tamoxifen [9, 10] and poorer long-term outcomes >5-10 years post-diagnosis [11–14]. Consistent with clinical observations, we identified tamoxifen-resistance in ER-positive ILC models, including *de novo* ER partial agonism by anti-estrogens [15, 16]. These data suggest that ER function and signaling is unique in the context of ILC.

Studies using laboratory models of ILC and tumor profiling through the Cancer Genome Atlas (TCGA) support that aspects of cell signaling are distinct in ILC versus IDC cells. TCGA analyses comparing Luminal A ILC to Luminal A IDC [4] using reverse-phase protein array (RPPA) data identified differential activity of PTEN and downstream Akt signaling. Additionally, each of the transcriptional subtypes of ILC (Reactive, Immune, and Proliferative) showed distinct signaling features in RPPA analyses (eg. high c-kit; high STAT5; high DNA repair protein signature; respectively). Consistent with distinct signaling contexts in ILC vs IDC, we reported that ER regulates unique target genes in ILC cells via distinct ER DNA binding patterns [15]. These ILC-specific ER target genes mediate ILC-specific signaling pathways that are critical for endocrine response and resistance, for example estrogen-driven atypical Wnt signaling via WNT4 [17, 18]. However, our understanding of ER-driven signaling at the protein level in ILC cells remains limited, as studies to date either cannot define dynamic changes caused by ER activation (ie. are from static samples as in TCGA) or are focused on the ER-driven transcriptome. Proteomic studies in ILC with estrogen or anti-estrogen treatment are needed to better understand dynamic ER-driven signaling in ILC.

We identified the Wnt ligand WNT4 as a critical signaling molecule induced by ER specifically in ILC cells [17]. WNT4 is unique among the Wnt protein family in its diverse cell type-specific roles, having been shown to either activate or suppress both canonical and non-canonical Wnt signaling pathways (discussed in [18]). In the normal mammary gland, WNT4 is induced by progesterone in progesterone receptor (PR) positive luminal epithelial cells, then secreted to act in a paracrine manner to activate canonical β-catenin-dependent signaling in neighboring myoepithelial cells [19–22]. In ILC cells, WNT4 regulation and signaling is ‘hijacked’ and falls under the direct control of ER [15, 17], but the mechanism by which WNT4 engages downstream signaling is unclear. Hallmark genetic loss of E-cadherin *(CDH1)* in ILC cells is associated with catenin protein dysfunction, including destabilization and loss of β-catenin protein in cell lines and tumors, resulting in impaired canonical Wnt signaling [17, 23]. Further, we recently reported that WNT4 secreted from ILC cells (and other cancer models) is dysfunctional in paracrine activity, instead activating signaling by a cell-autonomous, intracellular mechanism (ie. “atypical” Wnt signaling [18]). Taken together, defining ER-driven and WNT4-driven signaling is necessary to both understand ILC biology and identify new target treatments for ILC.

To address these gaps in our understanding of ER function in ILC, we used RPPA analyses of ILC models to characterize ER-driven signaling in ILC and to determine the role of WNT4 in mediating ER-driven signaling. These studies identified that estrogen activates distinct protein signaling pathways in ILC cells when compared to IDC cells, including a specific component of PI3K/Akt/mTOR signaling. Within the PI3K/Akt/mTOR signaling pathway, WNT4 was required most notably for downstream mTOR activity, in addition to regulation of mitochondrial function. These observations led us to further examine the mechanisms by which ER-driven WNT4 signaling mediates cell proliferation and survival, as well as endocrine response and resistance, in ILC cells.

## RESULTS

### Estrogen activates a distinct subset of PI3K-mTOR related signaling in ILC cells via WNT4

We profiled estrogen-driven signaling by RPPA, comparing hormone-deprived cells (vehicle treated, 0.01% EtOH) vs 1nM estradiol (E2) for 24hrs. RPPA was performed with ILC cell lines MDA MB 134VI (MM134) and SUM44PE (44PE) compared to IDC cell line MCF-7 (**Figure 1A**). Consistent with canonical pathways activated by ER, induction of ER target MYC, along with other cell cycle proteins, was observed in all three cell lines (**Figure 1B**). Other shared ER targets included activation of PI3K pathway proteins (eg. phospho-p70S6K, phospho-S6-S235/ S236) and suppression of cleaved caspase 7. We identified 18 proteins regulated by ER in MM134 and 44PE, but not in MCF-7 (ILC-specific ER targets, **Figure 1C**). These mainly represent either PI3K-related signaling (eg. phospho-S6, phospho-mTOR, total MCL-1) or transcriptional control (eg. NOTCH, SNAI1; we reported ER regulation of SNAI1/SNAIL previously [24]). Of note, estrogen suppressed total histone H3 in ILC, along with two H3 post-translational modifications (the latter reflective of the total H3 change). However, altering lysis conditions showed that H3 detected in RPPA represents soluble (non-nucleosomal) H3, rather than total cellular H3 (**Supplemental Figure 1A**, also see RPPA lysis conditions in Materials and Methods). Based on this, estrogen-driven decrease in soluble H3 is consistent with chromatin remodeling [25–28].

**Figure 1.**
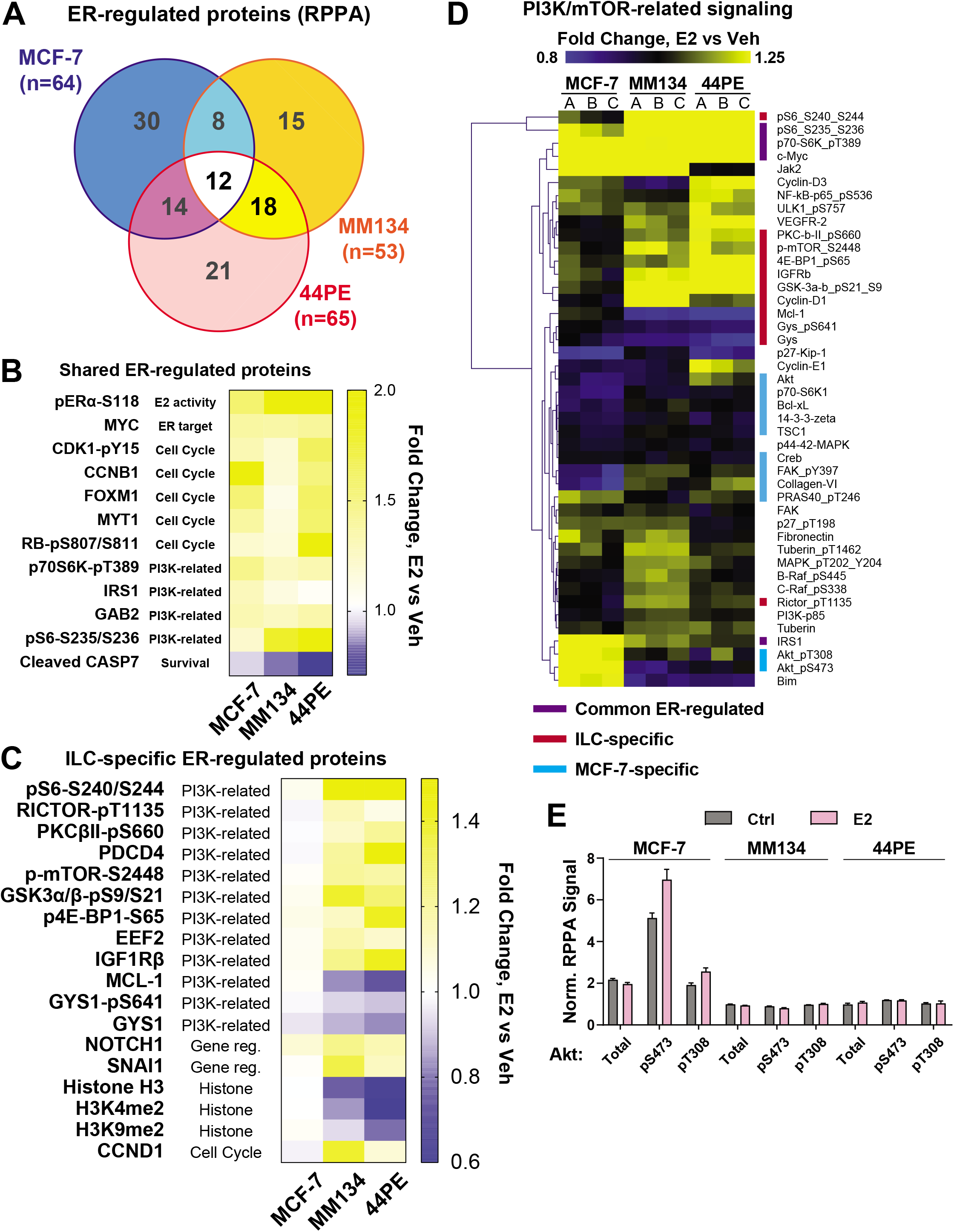
Estrogen regulates distinct protein targets in ILC cells versus MCF-7 cells. (A), Cells were hormone-deprived prior to treatment with 100pM E2 for 24hrs (biological triplicate; see Materials and Methods). RPPA protein signaling changes for 100pM E2 vs Veh shown (q<0.05 for MM134, 44PE; p<0.05 for MCF-7). (B-C), Heatmap for E2 vs Veh fold change (mean of triplicate samples). (D), PI3K pathway proteins were identified using KEGG pathways for PI3K/Akt/mTOR signaling. A/B/C represent biological triplicate samples. (E), Normalized Linear RPPA signal for total Akt or phospho-Akt (antibodies recognize AKT1/AKT2/AKT3).

Since ILC-specific E2-regulated targets were primarily related to PI3K-mTOR signaling, we examined all PI3K-related proteins from the RPPA that were changed by E2 treatment in any cell line to identify components of the signaling cascade in ILC (**Figure 1D**). Based on the observed estrogen-induced phosphorylation of mTOR, p70-S6K (RPS6KB1), and S6 in ILC cells (**Figure 1**), we expected to see an upstream increase in phosphorylated Akt [28]. However, estrogen-induced Akt phosphorylation was only observed in MCF-7 (**Figure 1D**), and levels of phospho- and total Akt were extremely low in ILC cells (**Figure 1E, Supplemental Figure 1B**). Similarly, Akt-mediated phosphorylation of PRAS40 was only estrogen-induced in MCF-7 suggesting mTOR is activated by another pathway in ILC.

Since non-canonical Wnt signaling has previously been linked to Akt-independent mTOR activation [29], we further investigated the role of WNT4 in estrogen-regulated signaling and mTOR activity in ILC cells. We compared hormone-deprived ILC cells treated with estrogen (as above) and transfected with non-targeting versus *WNT4*-targeting siRNA (siNT vs siWNT4, respectively). Estrogen-regulation of WNT4 and siWNT4 efficacy were confirmed by western blot (**Figure 2A**). Consistent with the critical role of WNT4 in ILC cell proliferation and survival [17], siWNT4 broadly dysregulated overall signaling (**Supplemental Figure 2A-B**). We compared siWNT4-mediated signaling changes to estrogen-regulated signaling in ILC cells and identified 9 targets for which estrogen-regulated changes were ablated by WNT4 knockdown (**Figure 2B**). WNT4 knockdown blocked estrogen-mediated FOXM1 induction, S6 phosphorylation, MCL-1 depletion, and soluble Histone H3 depletion in ILC cells; siWNT4 also increased phospho-ER (consistent with relieved negative feedback from mTOR signaling [30]). siWNT4 did not have similar effects on these targets in IDC cell lines MCF-7 or HCC1428 (**Figure 2C, Supplemental Figure 2C**). HCC1428 expresses high *WNT4* mRNA similar to that observed in ILC cell lines but does not depend on WNT4 for proliferation or survival [17], supporting that the identified signaling targets represent ILC-specific WNT4 signaling. Of note, we observed that the siNT construct caused a non-specific increase in phosphorylation of some mTOR targets (similar to our prior observations [17]), but this was blocked by siWNT4 (**Supplemental Figure 2D**). Based on the ILC-specific signaling targets of ER:WNT4, we hypothesized that WNT4 mediates ILC cell proliferation and survival via regulation of FOXM1 and mTOR.

**Figure 2.**
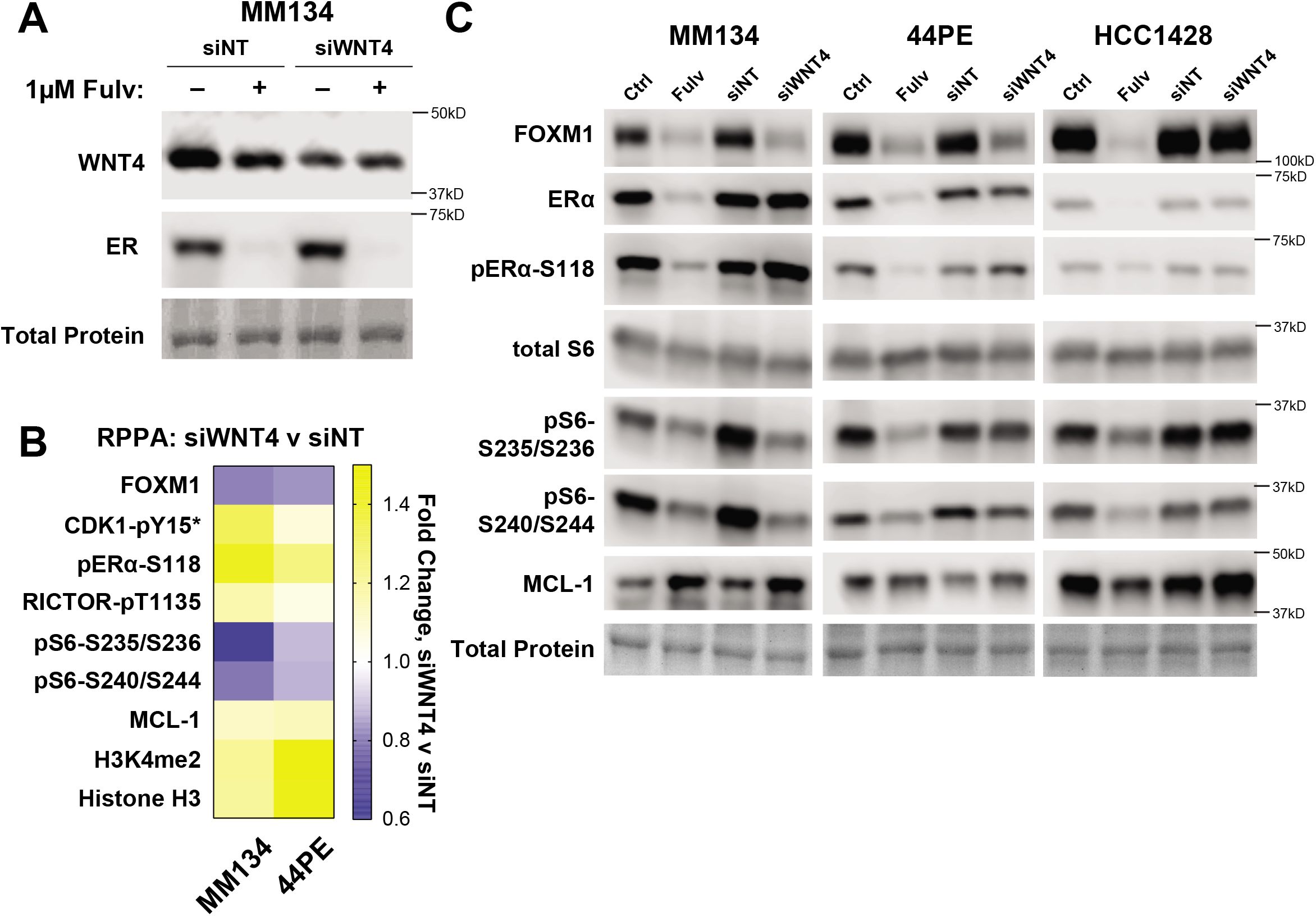
WNT4 is required for key components of ER signaling. (A), MM134 cells in full serum were reverse transfected with the indicated siRNA. 24hrs later, cells were treated with Fulvestrant, and lysates were harvested after an additional 24hrs. Total protein detected with Ponceau staining. (B), Cells were hormone-deprived prior to siRNA reverse transfection and subsequent treatment with 100pM E2 for 24hrs (biological triplicate; see Materials and Methods). RPPA protein signaling changes for siWNT4+E2 versus siNT+E2 shown (p<0.05) which overlapped with E2-driven signaling identified in Figure 1. *, siWNT4 effect is a false positive due to siNT-mediated suppression of signal verses E2/mock transfected (from Figure 1). (C), Cells were transfected and treated as in (A). Total protein detected with Ponceau staining.

### FOXM1 is not a direct target of WNT4 signaling

FOXM1 is a critical mediator of estrogen-driven cell cycle progression [31], has been linked to control of ER-driven transcription (in MCF-7 cells), and is a potential driver of anti-estrogen resistance [31–33]. This led us to examine whether FOXM1 is a critical WNT4 signaling target. Though *FOXM1* siRNA suppressed the proliferation of MM134 cells (**Supplemental Figure 3A**), this growth suppression was not equivalent to that observed with anti-estrogens or with *WNT4* siRNA. Whereas siWNT4 parallels treatment with fulvestrant and causes a G1 arrest, siFOXM1 causes a G2/M arrest (**Supplemental Figure 3B-C**), consistent with established roles of FOXM1 in progression through mitosis [34]. This lack of phenocopy between siWNT4/anti-estrogen and siFOXM1 suggested that decreased FOXM1 levels upon WNT4 knockdown may be indicative of knockdown-induced cell cycle arrest, rather than identifying FOXM1 as a direct WNT4 target. To confirm this, we examined whether CDKN1A/p21 knockdown, which partially restores proliferation after siWNT4 in ILC cells [17], would restore FOXM1 levels (ie. since p21 knockdown partially releases the G1 arrest, FOXM1 levels would increase despite WNT4 knockdown). In both MM134 and 44PE, siCDKN1A partially restored the FOXM1 decrease caused by siWNT4 (**Supplemental Figure 3D**), supporting that FOXM1 protein level is acting as a marker of cell cycle and G2/M progression. Consistent with a lack of a direct role in WNT4-driven cell cycle progression, we also did not observe any suppression of ER-driven transcription in MM134 by siFOXM1 (**Supplemental Figure 3E**). These data suggest that FOXM1 is not a direct target of ER and WNT4 in ILC but is an indirect target of ER/WNT4-driven cell cycle progression.

### WNT4 regulates mTOR signaling downstream of mTOR kinase activity

Our RPPA data showed estrogen treatment of ILC cells led to activation of mTOR signaling (eg. increased 4E-BP1, S6K/p70S6K, and S6 phosphorylation), but siWNT4 specifically suppressed S6 phosphorylation without altering phosphorylation of S6K (at T389) or 4E-BP1 (at S65). This suggests that WNT4 is required for mTOR signaling downstream of mTOR kinase activity. We examined this further by immunoblot using small molecule inhibitors of mTOR (everolimus) or S6K (PF-4708671, [35]) compared to targeting ER (fulvestrant) or WNT4 (siWNT4) (**Figure 3A**). As expected, everolimus suppressed phosphorylation of all targets tested, whereas PF specifically suppressed phosphorylation of S6, downstream of S6K. We confirmed that targeting ER reduced S6K phosphorylation at both T389 and T421/S424 (the latter not covered by RPPA), as well as downstream S6 phosphorylation. Consistent with our prior data, WNT4 knockdown suppressed S6 phosphorylation without affecting phosphorylation of mTOR or S6K (T389). However, we identified that siWNT4 suppressed S6K phosphorylation at T421/S424 (**Figure 3A**, blue box). This suggests WNT4 plays a specific role in releasing S6K from its pseudo-substrate inhibitory loop [36], increasing S6K activity and downstream S6 phosphorylation.

**Figure 3.**
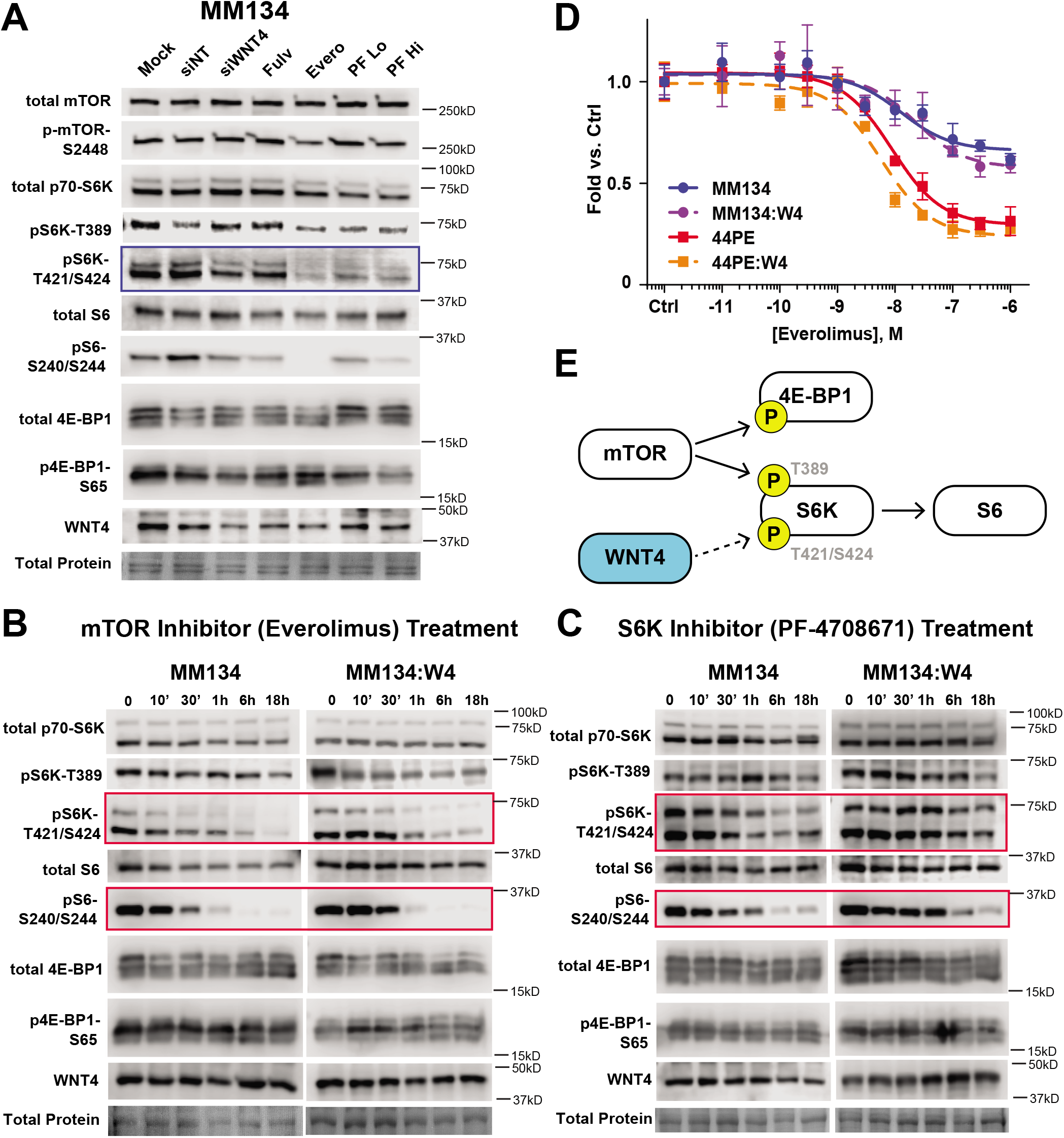
WNT4 independently controls S6 phosphorylation. (A), MM134 cells in full serum were either reverse transfected with the indicated siRNA for 72hrs, or treated with 100nM Fulvestrant, 10nM Everolimus, 10μM or 30μM PF-4708671 for 24hrs. (B-C), Cells in full serum were treated with 10nM Everolimus (B) or 30μM PF-4708671 (C) for the indicated time. Red boxes highlight time-dependent shifts in kinase inhibitor response driven by WNT4 over-expression (discussed in text). In (A-C), decreased 4E-BP1 phosphorylation is observed in downward size shift of total 4E-BP1; p-S65 results were inconsistent across studies. Total protein detected with Ponceau staining. (D), Cells were treated with increasing concentrations of Everolimus as indicated in biological replicates of 6 for 6d; total cell number was assessed by dsDNA quantification. Points represent mean of six biological replicates ±SD. (E), WNT4 controls phosphorylation of S6K at T421/S424, and as such required for S6K activity but dependent upon upstream S6K activation by mTOR.

Based on this observation, we hypothesized that WNT4 is necessary for, but unlikely sufficient for, S6K activity. We tested this by assessing the response to kinase inhibitors in WNT4 overexpressing MM134 cells. Upon treatment with everolimus, WNT4 over-expression modestly delayed the loss of S6K activity (pT421/S424 and S6 phosphorylation; **Figure 3B**, red boxes), but ultimately, downstream mTOR signaling was completely suppressed by everolimus within ~1h. Similar effects of WNT4 over-expression were observed in response to S6K inhibition, with prolonged rescue (**Figure 3C**, red boxes) but long-term S6K inhibition suppressed S6 phosphorylation. Consistent with this necessary but not sufficient role, WNT4 over-expression was unable to rescue inhibition of ILC cell proliferation by everolimus (**Figure 3D**). These data support that WNT4 is necessary for S6K function and mediates activity by driving T421/S424 phosphorylation. Upstream mTOR activity remains required for S6K activity, which similarly remains sensitive to direct small molecule inhibition (**Figure 3E**). A necessary-but-not-sufficient role for WNT4 in this pathway is consistent with similar characterization of WNT4 in the mammary gland [19, 37], and also suggests that other ER:WNT4 targets are critical mediators of proliferation and survival in ILC cells.

### ER:WNT4 regulation of MCL-1 is associated with metabolic dysregulation

In addition to regulating mTOR signaling in ILC cells, estrogen caused a decrease in levels of the BCL2-family protein MCL-1, which was reversed by WNT4 knockdown (**Figures 1–2**). ER regulation of MCL-1 likely occurs post-transcriptionally, as qPCR showed that *MCL1* gene expression is not regulated by estrogen in ILC cells (**Supplemental Figure 4A**). Translational regulation of MCL-1 has been previously linked to mTOR, with activation of mTOR signaling associated with increased MCL-1 levels [38]. However, our data show that estrogen activates mTOR while suppressing MCL-1, and targeting mTOR signaling (using everolimus or PF-4708671) does not affect MCL-1 levels (**Figure 4A**). These data are inconsistent with activated mTOR mediating increased MCL-1 translation. Thus ER:WNT4 regulation of MCL-1 in ILC cells is post-transcriptional but independent of mTOR signaling.

**Figure 4.**
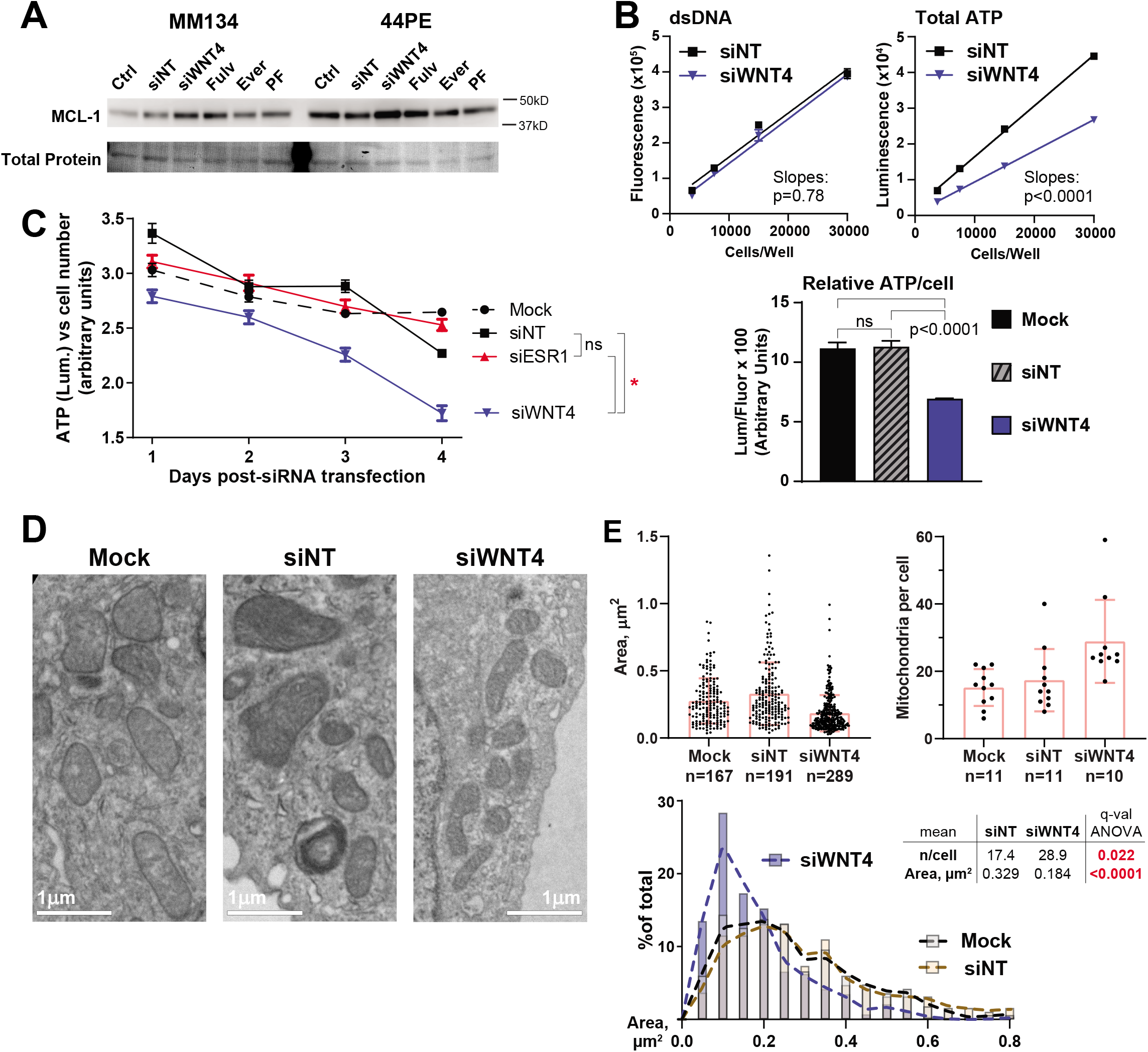
WNT4 knockdown compromises cellular metabolism and drives mitochondrial fragmentation. (A), Cells in full serum were either reverse transfected with the indicated siRNA for 48hrs, or treated with 100nM Fulvestrant, 100nM Everolimus, or 10μM PF-4708671 for 24hrs. Total protein detected with Ponceau staining. (B), MM134 cells were transfected with siRNA as indicated; 48h later cells were harvested and re-plated in a 2-fold serial dilution in technical quadruplicate. 24h later (72h post-transfection) dsDNA and total ATP were quantified from separate samples. Bar graph at bottom represents 30k/well data from standard curves (mock excluded from standard curves for clarity). Comparisons with ANOVA+Tukey correction. (C), MM134 cells transfected as indicated, with dsDNA and total ATP quantified at the indicated time from matched samples. Cell number was determined by interpolating dsDNA fluorescence vs a standard curve, and total ATP (luminescence) was normalized vs cell number. Comparisons by two-way ANOVA; α=0.01. *, p<0.0001. (D), MM134 cells were fixed for transmission electron microscopy 72h post-transfection. Representative images of mitochondrial morphology shown. (E), Mitochondrial dimensions were quantified from TEM images using ImageJ (see Materials and Methods). Box-and-whiskers represent mean ± SD. Histogram is derived from area data shown as dot plot at top.

Given the canonical role of MCL-1 as an anti-apoptotic BCL2-family protein [39], estrogen-driven reduction in MCL-1 levels contrasts the important roles of ER:WNT4 in mediating cell survival [17]. Based on this, we hypothesized that estrogen-mediated repression of MCL-1 may “prime” cells for apoptosis (e.g. to facilitate involution [40, 41]), but that anti-estrogens would make ILC cells resistant to apoptosis (ie. by inducing MCL-1). To test this, we treated ILC cells with BH3 mimetics (inhibitors of anti-apoptotic Bcl-2-family proteins) alone or in combination with fulvestrant and/or MCL-1-specific BH3 mimetic A-1210477. Though blocking MCL-1 modestly sensitized cells to other BH3 mimetics, fulvestrant caused no shifts in sensitivity to any tested combination of BH3 mimetics (**Supplemental Figure 4B**). This suggests that ER:WNT4 regulation of MCL-1 levels is not related to control of intrinsic apoptosis.

MCL-1 also plays a critical role in mitochondrial dynamics, regulating the balance of mitochondrial fission vs fusion and, subsequently, cellular capacity for oxidative phosphorylation [42–48]. Based on this, we hypothesized that ER:WNT4 control of MCL-1 indicates a role for WNT4 in mitochondrial dynamics and cellular metabolism in ILC cells. Consistent with this, siWNT4 caused a decrease in total ATP content per cell (**Figure 4B**). The decrease in cellular ATP levels was progressive over a 1-4d time course after siWNT4 (**Figure 4C**). ER knockdown did not produce a similar decrease in ATP levels, which parallels our prior observations that WNT4 knockdown, but not ER inhibition, induces cell death in ILC cells [17]. This suggests that a more profound depletion of WNT4 is required to effect mitochondrial function. Decreased ATP levels and increased MCL-1 after WNT4 knockdown is consistent with mitochondrial fission or fragmentation [45, 48], so we used transmission electron microscopy to investigate mitochondrial phenotype after WNT4 knockdown. At 72h post-transfection (ie. during progressive loss of ATP production, **Figure 4B-C**), cells with siWNT4 had an increased number of smaller mitochondria (**Figure 4D**). Cells with control siRNA had a mean of 17.4 mitochondria per cell with a mean area of 0.329μm^2^; siWNT4 caused a >65% increase in mitochondria per cell (mean = 28.9) while decreasing area by ~45% (mean = 0.184μm^2^) (**Figure 4E**). This supports that WNT4 knockdown drives mitochondrial fission or fragmentation. Taken together, these data suggest regulation of MCL-1 is related to a critical role for ER:WNT4 signaling in mitochondrial function and cellular metabolism.

### WNT4 signaling is re-activated during anti-estrogen resistance in ILC cells

We previously reported that long-term estrogen deprived (LTED; mimicking AI-resistance) variants of ILC cell lines remain dependent on WNT4 [17], and hypothesized that WNT4-driven pathways identified in the parental cells would also be active in LTED models. LTED models 44:LTED/A and 134:LTED/E (derived from 44PE and MM134, respectively [17]) were analyzed by RPPA and compared to hormone-deprived parental cells to identify signaling adaptations during LTED. LTED cells were also transfected with siNT or siWNT4 as above to identify WNT4-driven signaling during LTED.

The majority of protein changes in LTED vs parental comparisons were shared between models (**Figure 5A, Supplemental Figure 5A**). Of note, this included upregulation of FASN, confirming the transcriptional upregulation of *FASN* and lipid metabolism genes in these models that we recently reported (**Supplemental Figure 5B**) [49]. RPPA changes specific to 44:LTED/A (increased ER and OCT4) vs 134:LTED/E (decreased ER, increased phospho-NFκB) also confirmed model-specific features of LTED that are linked to WNT4 upregulation as we reported previously (**Supplemental Figure 5B**) [17]. Comparing protein changes in LTED models versus those caused by WNT4 knockdown identified 14 protein changes induced during LTED which are mediated by WNT4 signaling (**Figure 5B-D**). WNT4-driven signaling in LTED included key ER:WNT4 targets described above, eg. S6 phosphorylation and MCL-1 suppression, which were confirmed by immunoblotting (**Figure 5E**). These WNT4-driven signaling changes were also confirmed in additional ILC:LTED cell lines (**Supplemental Figure 6**). WNT4 regulation of downstream pathways, eg. cell cycle, mTOR signaling, and metabolism, remain active and critical to the LTED phenotype in ILC cells.

**Figure 5.**
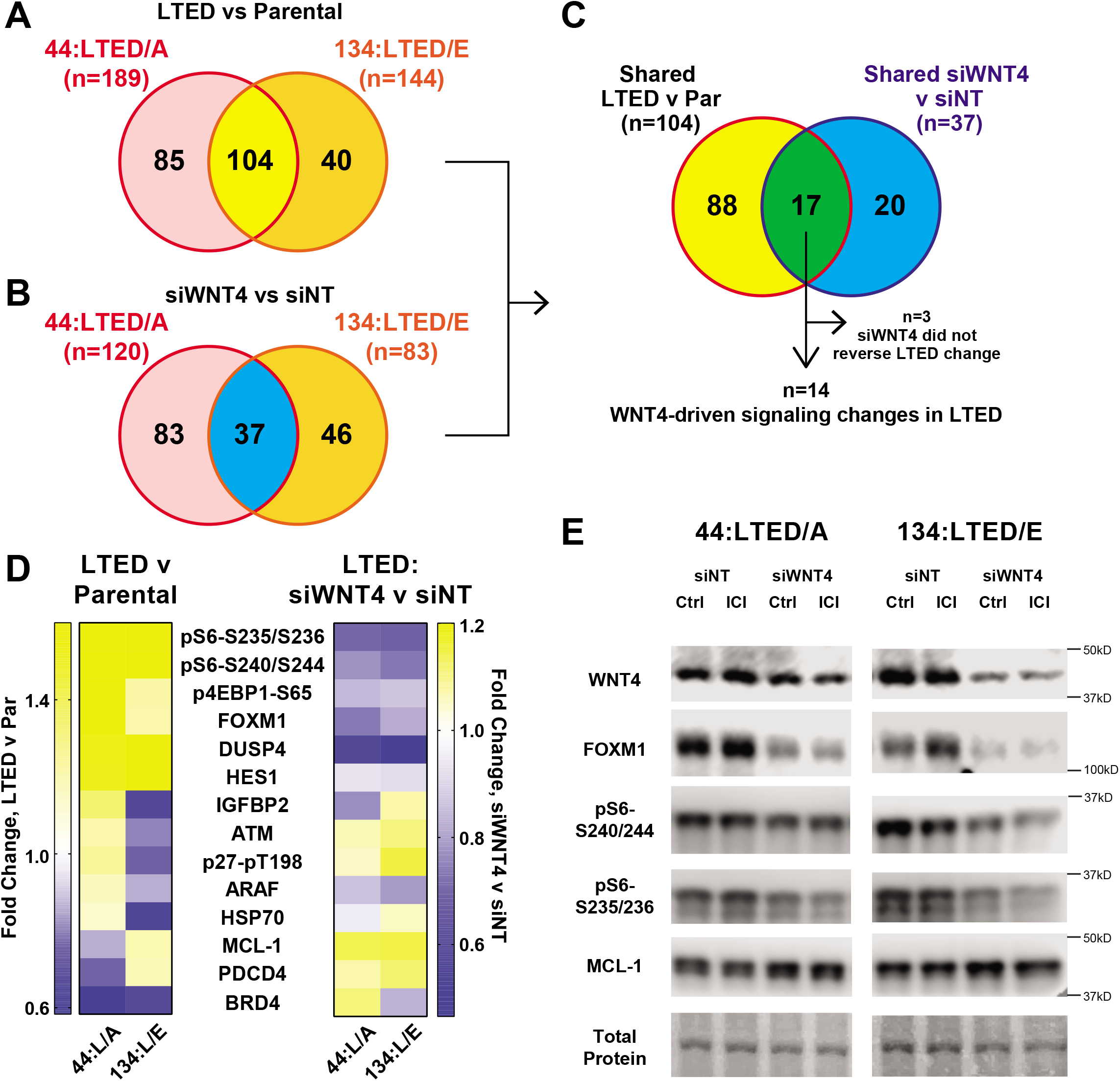
ER:WNT4 signaling targets are similarly active in anti-estrogen resistant ILC models. (A), LTED cells were compared to their hormone-deprived (vehicle treated) parental cell line; differences at q<0.05 shown. (B), LTED cells were reverse transfected with the indicated siRNA for 48hrs prior to harvest for RPPA analyses. RPPA protein signaling changes for siWNT4+E2 versus siNT+E2 shown (p<0.05). (C), Overlap between signaling activation in LTED (from (A)) versus WNT4-driven signaling (from (B)) to identify specific WNT4-driven signaling that is activated during anti-estrogen resistance. For n=3, siWNT4 did not reverse the changes seen in LTED v Parental in at least one of the two model systems. (D), Heatmap of n=14 genes from (C) for LTED v Parental fold change (left) and siWNT4 v siNT fold change (right). Boxes show mean of biological triplicate samples. (E), LTED cells (in hormone-deprivation) were reverse transfected with the indicated siRNA. 24hrs later, cells were treated with 1 μM Fulvestrant or vehicle, and lysates were harvested after an additional 24hrs. Total protein detected with Ponceau staining.

### Atypical WNT4 signaling functions in diverse tumor types

To determine whether similar WNT4-driven signaling functions in tumor tissues (including tumors beyond ILC), we explored data from the Cancer Genome Atlas (TCGA) using the cBio Portal [50]. We examined ILC and tumors related to tissues that require WNT4 in development (eg. kidney, adrenal, lung, ovary, uterus [51–54]), and identified tumors that over-expressed *WNT4* mRNA (**Figure 6A**). Comparing *WNT4* high versus other tumors, we looked for differences in protein signaling in RPPA data. Of note, many tumor types including ILC were limited by smaller sample sizes with associated RPPA data (eg. ILC, total n=160, high *WNT4* n=29). However, renal clear cell carcinoma (RCC; total n=515, high *WNT4* n=89), lung adenocarcinoma (LuA; total n=533, high *WNT4* n=57), and serous ovarian cancer (OvCa; total n=307, high *WNT4* n=42) presented differences in protein signaling by RPPA in *WNT4* high tumors (**Supplemental Figure 7A**). OvCa was unique in having an upregulation of PI3K/mTOR pathway phospho-proteins in high *WNT4* tumors (**Figure 6B, Supplemental Figure 7A-B**; MCL-1 data are not available in these datasets). Shared signaling changes were limited; differentially regulated proteins in all 3 tumors types (n=10 targets) were not similarly induced/repressed in all three tumor types, with the exception of high *WNT4* being associated with decreased ER levels (**Supplemental Figure 7C**). Changes in total β-catenin levels in high *WNT4* tumors were seen in LuA and RCC, but not OvCa (**Supplemental Figure 7D**). This suggests WNT4 in OvCa may activate non-canonical (β-catenin-independent) signaling that converges on mTOR as in ILC. Consistent with this, we previously identified that WNT4 engages atypical intracellular signaling in both ILC and OvCa cell lines [18]. Paralleling our observations in ILC cells, high *WNT4* in OvCa was associated with increased mTOR signaling activity, eg. phosphorylation of 4E-BP1, p70-S6K, and S6 (**Figure 6C**). Further, high *WNT4* expression in OvCa was associated with shorter overall survival (median OS, 28.5mo v 45.0mo; **Figure 6D**). Though RPPA data for ILC tumors were too limited to identify robust associations between *WNT4* expression and cell signaling, high *WNT4* was associated with decreased PTEN protein levels, consistent with activated downstream mTOR signaling (**Supplemental Figure 7E**). Additionally, high *WNT4* was associated with decreased phospho-Akt-S473, paralleling low phospho-Akt levels in our cell line data. Supporting this link between WNT4 and mTOR signaling in breast cancer, we found that in ER+ breast cancer, genes correlated with WNT4 expression were enriched for the Hallmark mTOR signaling signature (**Supplemental Figure 7F**). These data suggest that non-canonical WNT4 signaling via the mTOR pathway is critical in multiple tumor types, including ILC and OvCa.

**Figure 6.**
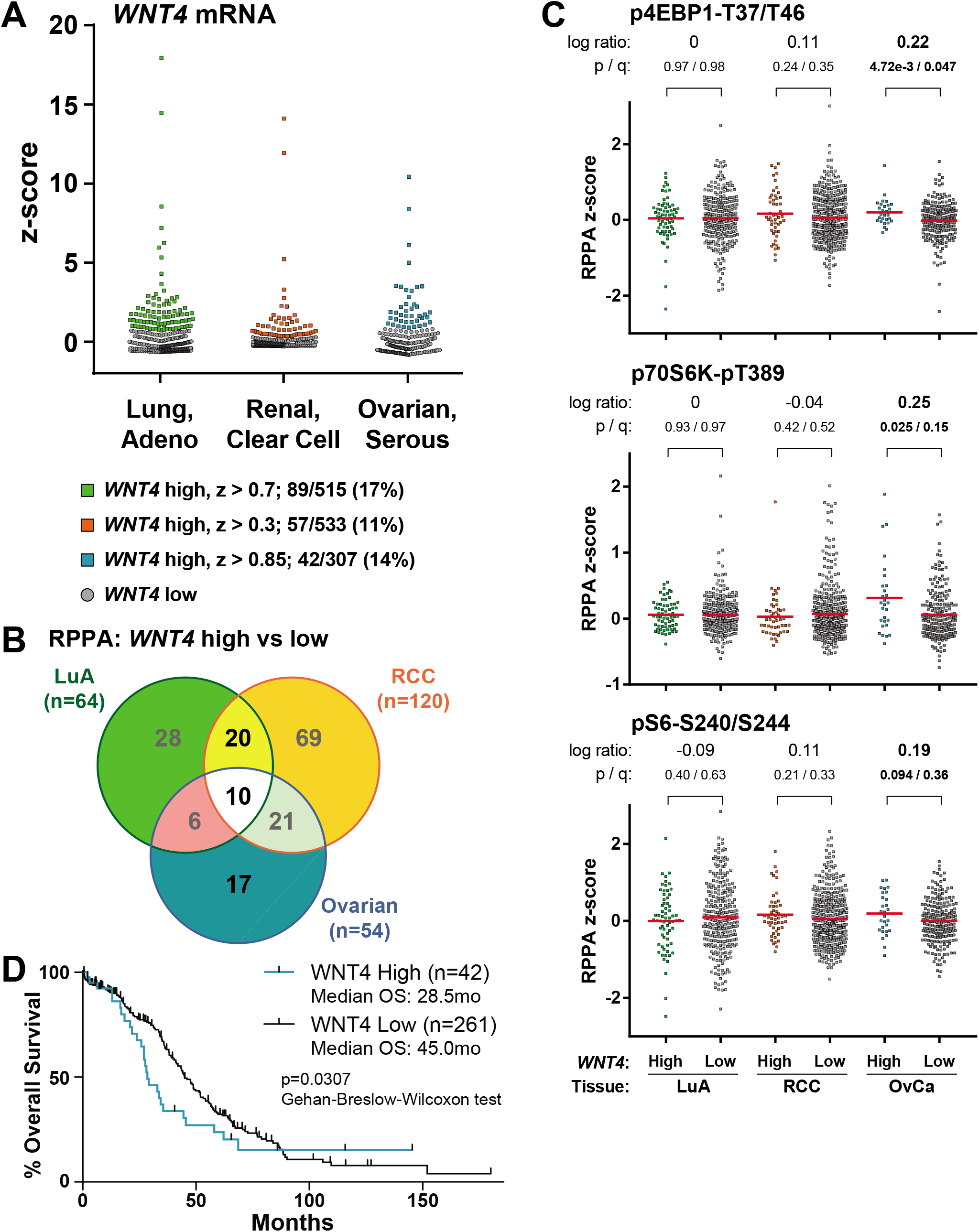
High *WNT4* expression is associated with activated mTOR signaling in serous ovarian cancer. (A), RNAseq z-scores used to identify tumors with high *WNT4* mRNA levels in the indicated tumor types from TCGA data. Points represent individual tumor samples. (B), cBio ‘Enrichment’ tool used to identify differences in RPPA signals (p<0.1) for high versus low *WNT4* OvCa samples. Colored v Gray points represent individual tumor samples with high/low *WNT4* from (A). Line represents mean RPPA signal for the indicate group. Log ratio shown as High v Low *WNT4*. (D), Overall survival data derived from cBio ‘Survival’ tool. High v Low *WNT4* as in (A).

## Discussion

Clinical and laboratory data support that ILC represents a unique context for estrogen receptor signaling, which may be mediated in part by ER-induced *WNT4* expression. The Wnt ligand WNT4 plays a critical role in the normal development of the mammary gland, and similarly is required for endocrine response and resistance in ILC cells [17]. However, WNT4 targets myriad tissue-specific pathways, β-catenin-dependent signaling is dysfunctional in ILC, and we recently reported that WNT4 has unique activity as an intracellular signaling molecule [4, 17, 18]. As such, the mechanisms by which WNT4 mediates proliferation and survival in ILC cells are unclear. To address this, we used RPPA analyses to profile ER-driven signaling in ILC cells and identify ER-driven signaling that requires WNT4 (ie. ER:WNT4 signaling). Our studies identified that in ILC cells, ER:WNT4 signaling mediates downstream activity of mTOR signaling via p70-S6K; WNT4 is required for p70-S6K phosphorylation (T421/S424) and downstream S6 phosphorylation. We also identified that ER:WNT4 signaling suppresses total MCL-1 levels, which was not associated with differential sensitivity to pro-apoptotic dugs, but instead with metabolic dysfunction and mitochondrial fragmentation upon WNT4 knockdown. Parallel signaling mediated by WNT4 was identified in anti-estrogen-resistant models of ILC and related signaling pathways were also linked to *WNT4* overexpression in serous ovarian cancer tumors. These data provide new insight in to a poorly understood Wnt signaling pathway, and identify signaling pathways downstream of atypical intracellular WNT4 signaling functions [18]. These downstream pathways may be targetable to inhibit cell proliferation and survival mediated by WNT4.

Three independent large-scale genomic analyses of ILC (the Cancer Genome Atlas; RATHER; Desmedt *et al* European cohort) describe activation of PI3K/Akt/mTOR signaling as a major feature of ILC, including increased pathway activity versus matched IDC tumors [4, 55, 56]. Pathway activation is associated with reduced PTEN protein levels in ILC, increased phospho-Akt, and increased phospho-p70-S6K (T389) [4]. These observations parallel our data identifying that ER drives downstream mTOR signaling in ILC cells. However, we observed that ILC cell lines MM134 and 44PE have low levels of phospho-Akt versus MCF-7 cells, suggesting WNT4 in ILC is an alternative pathway for the activation of mTOR signaling. In this case, estrogen-driven WNT4 may regulate mTOR in the context of low PTEN, as high *WNT4*-expressing ILC had decreased PTEN levels and decreased phospho-Akt (**Supplemental Figure 7E**). Notably, WNT4 was shown to be a driver of tumorigenesis in a PTEN-null model of epithelial ovarian cancer, and in this model WNT4-driven invasion/migration was insensitive to Akt inhibition [57]. Taken together with our observations, WNT4 likely regulates mTOR signaling downstream of Akt. Though WNT4 itself has not been mechanistically linked directly to the mTOR pathway, β-catenin-independent Wnt signaling was shown by Inoki et al to regulate mTOR activity via GSK3, independent of Akt phosphorylation [29]. GSK3 phosphorylation was strongly E2-induced in ILC cells (**Figure 1**), suggesting GSK3 activity links WNT4 to mTOR. Similarly, GSK3 phosphorylation was among the most strongly increased signaling events in high *WNT4*-expressing OvCa (**Supplemental Figure 7B**). However, GSK3 phosphorylation was not affected by WNT4 knockdown in ILC cells by RPPA (**Figure 2**, **Supplemental Figure 2**). Further, Inoki et al showed that Wnt signaling modulated p70-S6K phosphorylation at T389. We observed that while estrogen did increase phosphorylation at T389, WNT4 knockdown had no effect on T389 phosphorylation. WNT4 instead regulates phosphorylation at T421/S424 (**Figure 3**). T421/S424 are part of a cluster of phosphorylation sites in the C-terminal auto-inhibitory domain of p70-S6K (via pseudo-substrate inhibition), and their phosphorylation is necessary for S6K activation [36]. The regulation of phosphorylation at these sites is not fully understood, but kinases including ERK1/2 [58, 59], CDK5 [60, 61], JNK [62], and TBK1 [63] have been shown to target these sites. Neither JNK nor ERK1/2 were regulated by WNT4 knockdown in our RPPA, and we showed that JNK is not a target of non-canonical paracrine WNT4 signaling in MM134 cells [18]. CDK5 may be a potential link between WNT4 and p70-S6K, as CDK5 has been linked to non-canonical Wnt signaling [64] as well as mitochondrial function in breast cancer cells [65]. *CDK5* gene expression is also positively correlated with *WNT4* expression in TCGA in ER+ breast cancer (Spearman ρ = 0.14; q = 0.00018) and specifically ER+ ILC (Spearman ρ = 0.17; q = 0.028)[50]. Though CDK5 or other kinases (or phosphatases) may represent key steps linking WNT4 downstream to p70-S6K, defining the mechanism by which WNT4 initiates and propagates signaling is an important future direction.

Though our data suggest WNT4 signaling activates mTOR/p70-S6K downstream of Akt, a recent study by Teo et al linked loss of E-cadherin (ie. the hallmark feature of ILC [4]) to Akt activation and signaling. Notably, this link was identified using ER-negative ILC models (murine p53/CDH1-null lines and IPH-926)[66], suggesting that across ILC, subsets of cells/tumors exist with distinct modes of PI3K/Akt/mTOR signaling. Supporting this, among Luminal A ILC, TCGA RPPA identified differential levels and phosphorylation of PI3K/Akt/mTOR pathway proteins in the mRNA subtypes (eg. low p70S6K and Raptor in Reactive; high phospho-PRAS40 and phospho-mTOR in immune; Proliferative lacked distinct differences in this pathway) [4]. Another important context for PI3K-pathway signaling in ILC is likely the tumor microenvironment and metastasis. Tasdemir et al recently reported that in ultra-low attachment conditions (ie. requiring anchorage-independence, mimicking metastasis), ILC cell lines were uniquely able to sustain PI3K-pathway activation versus IDC cell lines [67]. Further mechanistic studies are needed to link biomarkers with PI3K/Akt/mTOR signaling activity to identify precision targets for therapy, ie. the ideal point on this pathway to intervene in an individual tumor.

ER:WNT4 regulation of MCL-1 and mitochondrial function suggest that WNT4 is critical for cellular metabolism in ILC cells. We observed that WNT4 knockdown leads to impaired ATP production and mitochondrial fragmentation (**Figure 4**), and both of these phenotypes precede the induction of cell death (~4d post-siWNT4 transfection, [17]). This suggests that WNT4-mediated control of metabolism and/or mitochondrial function are critical to ILC cell survival. Wnt signaling (both β-catenin-dependent and -independent signaling) has been previously linked to these processes [68–70], but direct roles for WNT4 in mitochondrial function have not previously described. However, one recent study demonstrated that Wnt4 overexpression could rescue a defect in mitochondrial function and dynamics caused by deletion of PTEN-inducible kinase 1 (PINK1) in Drosophila [71]. Though the mechanism of rescue is unclear, Wnt4 overexpression rescued flight defects in the PINK1 knockout flies, increasing ATP production and restoring mitochondria membrane potential in flight muscles. Future studies will further examine whether WNT4 plays a similar role in cancer cells. Further, key questions raised by our study include: 1) Does WNT4 directly regulate MCL-1 (levels, localization, etc [42, 45]) or indirectly via the mitochondria; 2) Is mitochondrial fragmentation on WNT4 knockdown due to WNT4 control of mitochondrial fission/fusion machinery, or control of biogenesis vs mitophagy; 3) Does intracellular WNT4 localize to or act directly at the mitochondria? In considering targeting metabolic functions regulated by WNT4, defects caused by WNT4 knockdown may also be caused by suppressing *WNT4* expression using anti-estrogens or WNT4 levels using mTOR pathway inhibitors. Targeting therapy-induced metabolic vulnerability may be a powerful combination treatment approach for WNT4-driven cancers.

The similar signaling pathways associated with WNT4 in ILC and OvCa may parallel the critical role of WNT4 in both tissues of origin, as well as related tumor biology. In addition to being required for mammary gland development, WNT4 is required for the development of the ovary and Mullerian tissues [51, 72], female sex differentiation [72, 73], and fertility [52, 74]. Accordingly, WNT4 dysfunction is linked to a range of endocrine and gynecologic pathologies, including endometriosis [75], uterine fibroids [76], and ovarian cancer [77]. For ovarian cancer that originates from fallopian tube epithelium (FTE), transformed FTE cells must migrate to and invade the ovary to establish a tumor. As noted above, WNT4 is required for cell migration and ovary invasion in murine PTEN-null models of FTE-derived ovarian cancer [57], suggesting WNT4 is critical in early ovarian tumorigenesis. Perhaps consistent with this role of WNT4 in migration/invasion, both OvCa and ILC metastasize to the abdomen/peritoneal cavity, ILC being unique among breast cancers in this regard [5, 6]. Defining WNT4 signaling in these tumors may provide new insight in to targeting metastatic OvCa and ILC.

WNT4 signaling is a key mediator of endocrine response and resistance in ILC, but given the dysfunction of β-catenin-dependent signaling in ILC and novel functions of WNT4 as an intracellular signaling molecule, WNT4 signaling pathways in ILC must be identified. In this study, our RPPA analyses identified mTOR signaling and MCL-1/mitochondrial function as two key downstream effectors of WNT4 in ILC. Parallel pathways associated with WNT4 were also identified in serous ovarian cancer, suggesting that WNT4 signaling is important in multiple tumor types. Future studies will determine how WNT4 is mechanistically linked to these targets, including identification of the WNT4 “receptor” in ILC cells. Furthering our understanding of WNT4 signaling will improve opportunities for precision treatment approaches by targeting WNT4 signaling for patients with ILC, or OvCa, and support the development of strategies to overcome anti-estrogen resistance for patients with ILC.

## Materials and Methods

### Cell culture

MDA MB 134VI (MM134) and SUM44PE (44PE) were maintained as described [15]. MCF-7 and HCC1428 were maintained in DMEM/F12 (Corning Life Sciences, cat#10092CV) supplemented with 10% fetal bovine serum (FBS; Nucleus Biologics, cat#FBS1824). Hormone-deprivation was performed as described [78] in IMEM (Gibco/ThermoFisher, cat#A10488) supplemented with 10% charcoal-stripped fetal bovine serum (CSS; prepared as described [78] with the same FBS as above). WNT4overexpressing models were previously described [18], and cultured in the same conditions as parental cell lines. Long-term estrogen deprived (LTED) model establishment and culture conditions were previously described [17]. All lines were incubated at 37°C in 5% CO_2_. Cell lines are authenticated annually via the University of Arizona Genetics Core cell line authentication service and confirmed to be mycoplasma negative every four months. Authenticated cells were in continuous culture <6 months.

17β-Estradiol (E2; cat#2824) and fulvestrant (fulv / ICI182,780; cat#1047) were obtained from Tocris Bioscience and dissolved in ethanol. Everolimus (evero; cat#11597), PF-4708671 (PF; cat#15018), ABT199 (cat#16233), ABT263 (cat#11500), WEHI539 (WEHI; cat#21478), and A1210477 (A121; cat#21113) were obtained from Cayman Chemical and dissolved in DMSO.

### RNA interference

siRNAs were reverse transfected using RNAiMAX (ThermoFisher) according to the manufacturer’s instructions. All constructs are siGENOME SMARTpool siRNAs (Dharmacon/Horizon Discovery): Nontargeting pool #2 (D-001206-14-05), Human *WNT4* (M-008659-03-0005), Human *FOXM1* (M-00976200-0005), Human *MCL1* (M-004501-08-0005), Human *CDKN1A* (M-003471-00-0005). Details regarding validation of the specific effects of the *WNT4* siRNA pool are previously described [17].

### Reverse phase protein array (RPPA)

Cells were hormone-deprived prior to reverse transfection with 10nM siRNA as described above. 24 hours post-transfection, cells were treated with vehicle (0.01% EtOH) or 100pM E2 and harvested 24 hours later (48h post-transfection; 24h post-treatment). Cells were lysed according to core facility instructions (see below, [79]). Briefly, cells were washed twice with cold PBS, and incubated on ice with 60μL of RPPA lysis buffer (see below) for 20’. Lysates were collected by scraping and centrifuged at ~16,000xg at 4°C for 10’. Supernatant was collected and utilized for RPPA analyses. RPPA lysis buffer was made with 1% Triton X-100, 50mM HEPES (pH 7.4), 150mM NaCl, 1.5mM MgCl2, 1mM EGTA, and 10% glycerol, supplemented with Halt Protease and Phosphatase Inhibitor Cocktail (Pierce/ThermoFisher, cat#78440).

RPPA analyses and data normalization were carried out by the MD Anderson Cancer Center Functional Proteomics Core Facility (August 2016; platform included 305 antibodies listed in **Supplemental Table 1**). Details for RPPA signal normalization and quality control provided by the core facility are in **Supplemental Document 1**; Normalized Linear data are presented in all analyses herein. Multiple testing correction was applied using the Benjamini–Hochberg method. Raw/normalized RPPA data and treatment comparisons will be deposited to an OSF page upon publication [1], or will be provided upon request.

### Immunoblotting

Whole-cell lysates were obtained by incubating cells in RPPA lysis buffer (above) for 30’ on ice. Cells were centrifuged at ~16,000xg for 15m at 4°C and the resulting supernatant was collected for analysis. Protein concentrations were measured and normalized using the Pierce BCA Protein Assay Kit (#23225). Protein loading was kept consistent by mass across matched experiments, and standard methods were used to perform SDS-PAGE. Proteins were transferred onto PVDF membranes. Antibodies were used according to manufacturer’s recommendations: WNT4 (R&D, MAB4751, cat# 55025, also see specific immunoblot considerations [18]); ERα (Leica 6F11, cat# ER-6F11-L-F); pER-S118 (Cell Signaling, 16J4, cat#2511); FOXM1 (Cell Signaling, D12D5, cat#5436); total S6 (Cell Signaling, 54D2, cat#2317); pS6-S235/236 (Cell Signaling, cat#2211); pS6-S240/244 (Cell Signaling, cat#2215); p70-S6K (Cell Signaling, 49D7, cat#2708); p-p70-S6K-T389 (Cell Signaling, 108D2, cat#9234); p-p70-S6K-T421/S424 (Cell Signaling, cat#9204); total mTOR (Cell Signaling, 7C10, cat#2983); p-mTOR-S2448 (Cell Signaling, D9C2, cat#5536); MCL-1 (Cell Signaling, D35A5, cat#5453); 4E-BP1 (Cell Signaling, cat#9452); p4E-BP1-S65 (Cell Signaling, 174A9, cat#9456); Histone H3 (Abcam, cat#ab1791). Secondary antibodies were used according to manufacturer’s instruction and were obtained from Jackson ImmunoResearch Laboratories (West Grove, PA, USA): Goat Anti-Mouse IgG (cat # 115-035-068), Goat Anti-Rabbit IgG (cat# 111-035-045) and Goat Anti-Rat IgG (cat# 112-035-062). Chemiluminescence was used to detect antibodies and either film or the LICOR c-Digit (LI-COR Biosciences, Lincoln, NE, USA) was used to develop the immunoblots. Total protein (Ponceau) served as a loading control. Representative immunoblots with size markers indicated are shown as **Supplemental Figure 8.**

### Cell proliferation and cell cycle analyses

Total double-stranded DNA was measured as a proxy for total cell number by hypotonic lysis of cells in ultra-pure H_2_O, followed by addition of Hoechst 33258 (ThermoFisher Scientific, #62249) at 1μg/mL in Tris-NaCl buffer (10mM Tris, 2M NaCl; pH 7.4) at equivalent volume to lysate. Fluorescence (360nm ex / 460nm em) was measured on a Bio-Tek Synergy 2 microplate reader. For cell cycle analyses, cells were fixed in 70% EtOH overnight at 4°C. After fixation, cells were treated with RNase A, and stained with 50μg/mL propidium iodide + 0.1% Triton X-100 in PBS overnight at 4°C. PI intensity was assessed on a Gallios Flow Cytometer (Beckman Coulter) at the U. Colorado Cancer Center Flow Cytometry Core Facility; at least 10,000 events were counted per sample.

### Quantitative PCR analyses

RNA extractions were performed using the RNeasy Mini kit (Qiagen); mRNA was converted to cDNA on an Eppendorf Mastercycler Pro (Eppendorf) and using Promega reagents: Oligo (dT)15 primer (cat# C110A), Random Primers (cat# C118A), GoScript 5x Reaction Buffer (cat# A500D), 25mM MgCl2 (cat# A351H), 10mM dNTPs (cat# U1511), RNasin Plus RNase Inhibitor (cat# N261B) and GoScript Reverse Transcriptase (cat# A501D). qPCR reactions were performed with PowerUp SYBR Green Master Mix (Life Technologies, cat #100029284) on a QuantStudio 6 Flex Real-Time PCR system (ThermoFisher). Expression data were normalized to *RPLP0*. Primer sequences were published previously [15, 17].

### Metabolic analyses

For analysis of ATP per cell, cells were reverse transfected as above, and at the indicated timepoint, parallel plates were assessed for total dsDNA (above) and total ATP (Cell Titer-Glo, Promega). Cell number by each method was normalized to a standard curve, and analyzed as a ratio of (Cell Number by ATP)/(Cell Number by dsDNA).

### Transmission electron microscopy and mitochondrial phenotype analysis

MM134 cells were reverse transfected with siRNA as above, and 48h later plated in technical duplicate per condition to 6cm plates. 24h later (ie. 72h post-transfection), cells were collected by saline-EDTA wash and scraping, then fixed and pelleted in 2% glutaraldehyde; further processing was performed by the U. Colorado Anschutz Electron Microscopy Center. Pellets were re-suspended in 5% agarose that was allowed to harden, and then cut into small pieces for processing. Embedded cells were first rinsed three times in 0.1 M Sodium Cacodylate buffer (pH 7.4), then post-fixed in 1% osmium tetroxide for 1 hour. After three washes in water, the cells were stained *en bloc* with 2% uranyl acetate for 1 hour at 4°C. Following dehydration through a graded ethanol series (50%, 75%, 95%, 100%) and propylene oxide for 10 minutes each, the samples were infiltrated with LX112 resin. The samples were embedded and cured for 48 hours at 60°C in an oven. Ultra-thin sections (60nm) were cut on a Reichert Ultracut S from a small trapezoid positioned over the cells, and were picked up on EMS copper mesh grids. Sections were imaged on a FEI Tecnai G2 transmission electron microscope (Hillsboro, OR) with an AMT digital camera (Woburn, MA). From collected images, mitochondrial dimensions were measured in ImageJ [80] as described by Tobias et al [81]. Briefly, mitochondrial outer membranes were traced using the Freehand Selection Tool, and then fit to ellipses. Area measurements were taken from the fit ellipses; data from the technical duplicates were combined.

### TCGA data analyses

TCGA data were accessed via the cBio portal in March-June 2019; provisional datasets were used for all analyses described. Z-score cutoffs for *WNT4* expression (RNAseq) were selected based on the minimum z-score needed to exclude any low *WNT4*-expressing tumors. All statistical analyses were derived from cBio analysis tools, ie. Enrichments>Protein>RPPA.

## Data availability

Data associated with experiments herein will be available at an Open Science Framework repository [1] (https://doi.org/10.17605/OSF.IO/7X8NG) upon publication, or upon request.

## Acknowledgements

We thank Dr. Jennifer Bourne and the CU Anschutz Electron Microscopy Center for their support and technical assistance.

