## Supplemental Figures 1-8 for "Estrogen controls mTOR signaling and mitochondrial function via WNT4 in lobular carcinoma cells"

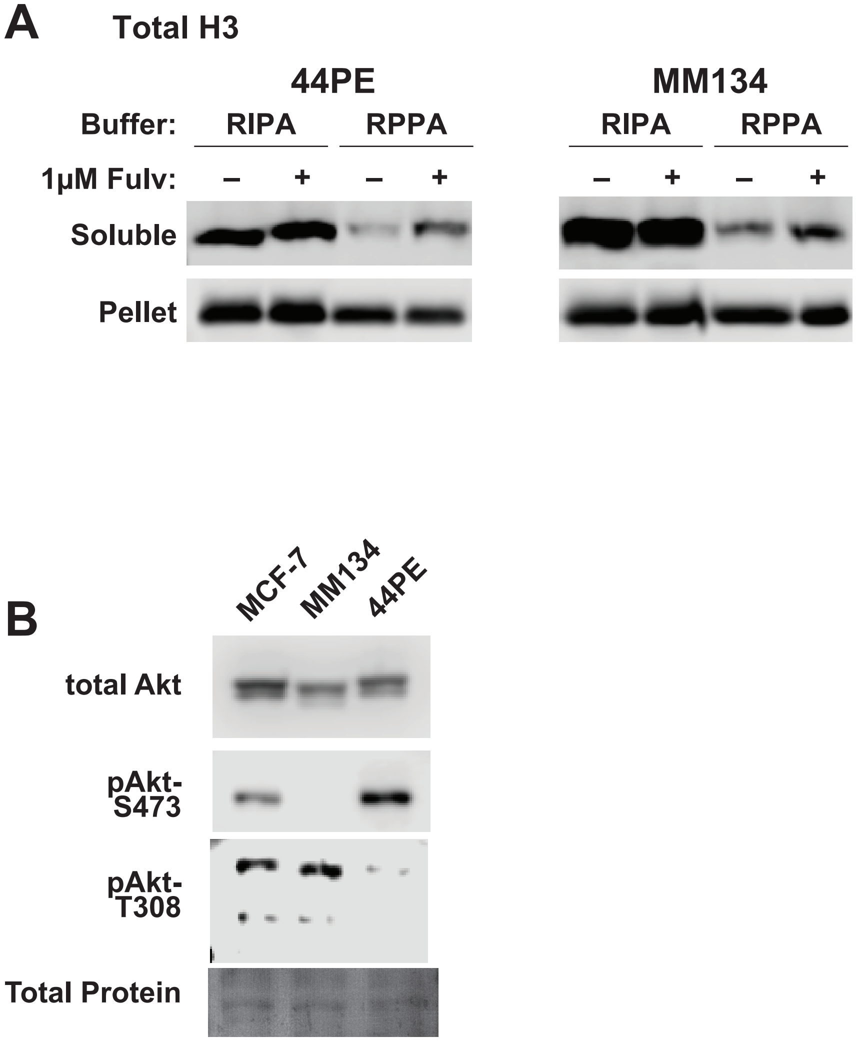


**Supplemental Figure 1. RPPA signal represents soluble Histone H3 and Akt/p-Akt pools.** (A), ILC cells were treated with Fulvestrant or vehicle for 24hrs and lysed with either RPPA buffer (as described in Materials and Methods) or RIPA buffer. RPPA buffer (lacks SDS) does not solubilize compacted chromatin, which is pelleted and removed prior to further processing (ie. pellet sample, which contains most of the cellular pool of histones). (B), Confluent (~80%) cells in full serum were lysed with RPPA buffer (as described in Materials and Methods). Total protein detected with Ponceau staining.

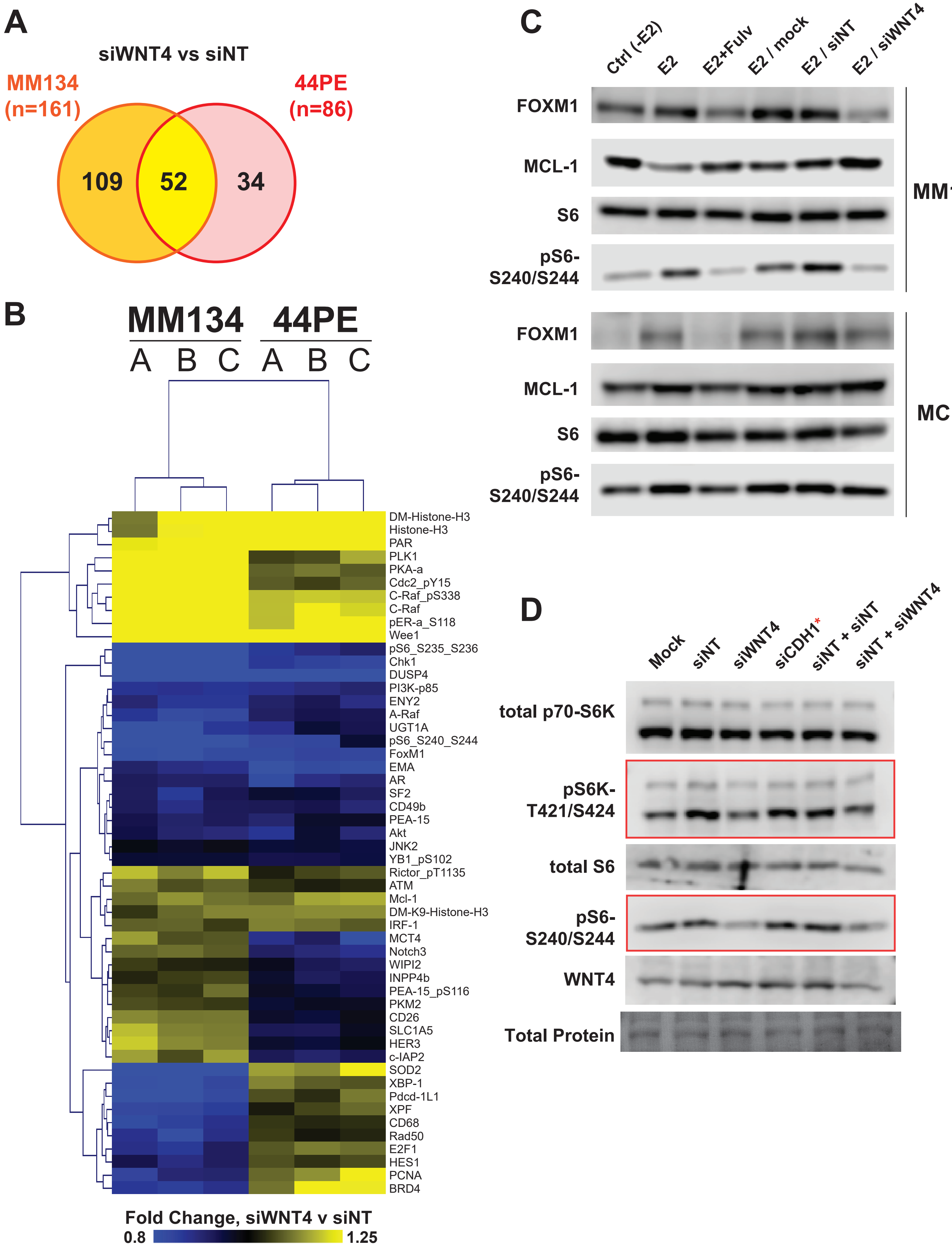

A
siWNT4 vs siNT
MM134
(n=161)
44PE
(n=86)
109
52
34
B
MM134
44PE
A B C A B C
DM-Histone-H3
Histone-H3
PAR
PLK1
PKA-a
Cdc2_pY15
C-Raf_pS338
C-Raf
pER-a_S118
Wee1
pS6_S235_S236
Chk1
DUSP4
PI3K-p85
ENY2
A-Raf
UGT1A
pS6_S240_S244
FoxM1
EMA
AR
SF2
CD49b
PEA-15
Akt
JNK2
YB1_pS102
Rictor_pT1135
ATM
Mcl-1
DM-K9-Histone-H3
IRF-1
MCT4
Notch3
WIPI2
INPP4b
PEA-15_pS116
PKM2
CD26
SLC1A5
HER3
c-IAP2
SOD2
XBP-1
Pdcd-1L1
XPF
CD68
Rad50
E2F1
HES1
PCNA
BRD4
Fold Change, siWNT4 v siNT
0.8
1.25
C
Ctrl (-E2)
E2
E2+Fulv
E2 / mock
E2 / siNT
E2 / siWNT4
FOXM1
MCL-1
S6
pS6-
S240/S244
MM134
FOXM1
MCL-1
S6
pS6-
S240/S244
MCF7
D
Mock
siNT
siWNT4
siCDH1*
siNT + siNT
siNT + siWNT4
total p70-S6K
pS6K-
T421/S424
total S6
pS6-
S240/S244
WNT4
Total Protein

**Supplemental Figure 2. WNT4 knockdown broadly effects signaling in ILC cells.** (A), Cells were hormone-deprived prior to siRNA reverse transfection and subsequent treatment with 100pM E2 for 24hrs (biological triplicate; see Materials and Methods). RPPA protein signaling changes for siWNT4+E2 versus siNT+E2 shown ($p<0.05$). (B), Heatmap for shared targets, siWNT4 vs siNT. A/B/C represent biological replicate samples. (C), Cells were treated and transfected as in (A). (D), MM134 cells in full serum were reverse transfected with the indicated siRNA for 72hrs. siCDH1 (targeting E-cadherin) was used as a pseudo-non-targeting construct, since ILC cells are negative for E-cadherin protein. The red boxes highlight where both siNT and siCDH1 increase mTOR target phosphorylation, suggesting a class effect of siRNAs. Combining siWNT4 with siNT reverses these inductions, confirming that WNT4 knockdown suppresses mTOR signaling. Total protein detected with Ponceau staining.

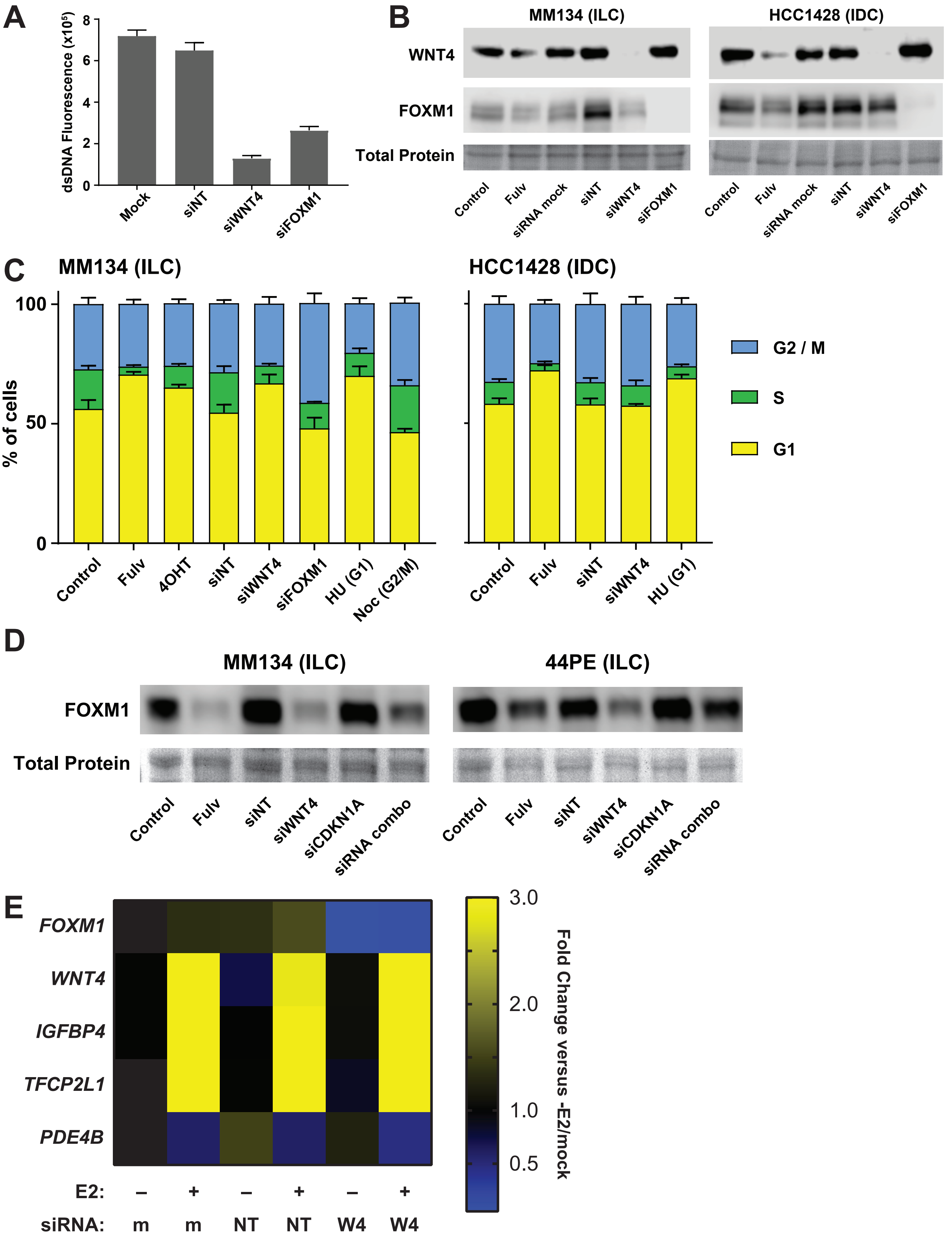
A
dsDNA Fluorescence (x10⁵)
8
6
4
2
0
Mock
siNT
siWNT4
siFOXM1
B
MM134 (ILC)
HCC1428 (IDC)
WNT4
FOXM1
Total Protein
Control
Fulv
siRNA mock
siNT
siWNT4
siFOXM1
C
MM134 (ILC)
HCC1428 (IDC)
% of cells
100
50
0
G2 / M
S
G1
Control
Fulv
4OHT
siNT
siWNT4
siFOXM1
HU (G1)
Noc (G2/M)
D
MM134 (ILC)
44PE (ILC)
FOXM1
Total Protein
Control
Fulv
siNT
siWNT4
siCDKN1A
siRNA combo
E
FOXM1
WNT4
IGFBP4
TFCP2L1
PDE4B
E2: – + – + – +
siRNA: m m NT NT W4 W4
3.0
2.0
1.0
0.5
Fold Change versus -E2/mock

**Supplemental Figure 3. FOXM1 is not a direct target of WNT4 signaling.** (A), MM134 cells in full serum were reverse transfected with the indicated siRNA, and proliferation was assessed by dsDNA quantification after 6d. Bars represent mean of 6 biological replicates ±SD. (B), Confirmation of WNT4 or FOXM1 knockdown. MM134 cells in full serum were reverse tranfected with the indicated siRNA. 24hrs later, cells were treated with Fulvestrant, and lysates were harvested an additional 24hrs later. Total protein detected with Ponceau staining. (C), Cells in full serum were transfected and treated as in (B) using 1μM Fulvestrant, 1μM 4-hydroxytamoxifen, 2mM hydroxyurea, or 50ng/mL nodocazole. Bars represent mean cell fraction per cell cycle phase of 3 biological replicate experiments ±SD. (D), Cells were transfected and treated as in (B). In siRNA combinations, siNT were used to increase total siRNA to 20nM in single siRNA samples. Total protein detected with Ponceau staining. (E), Hormone-deprived MM134 were reversed transfected with the indicated siRNA. 24hrs later, cells were treated with 100pM E2 or vehicle, and RNA was harvested after an additional 24hrs. Expression was normalized to *RPLP0*. Box represents mean fold change of biological triplicate samples, versus the -E2/mock control sample (Column 1).

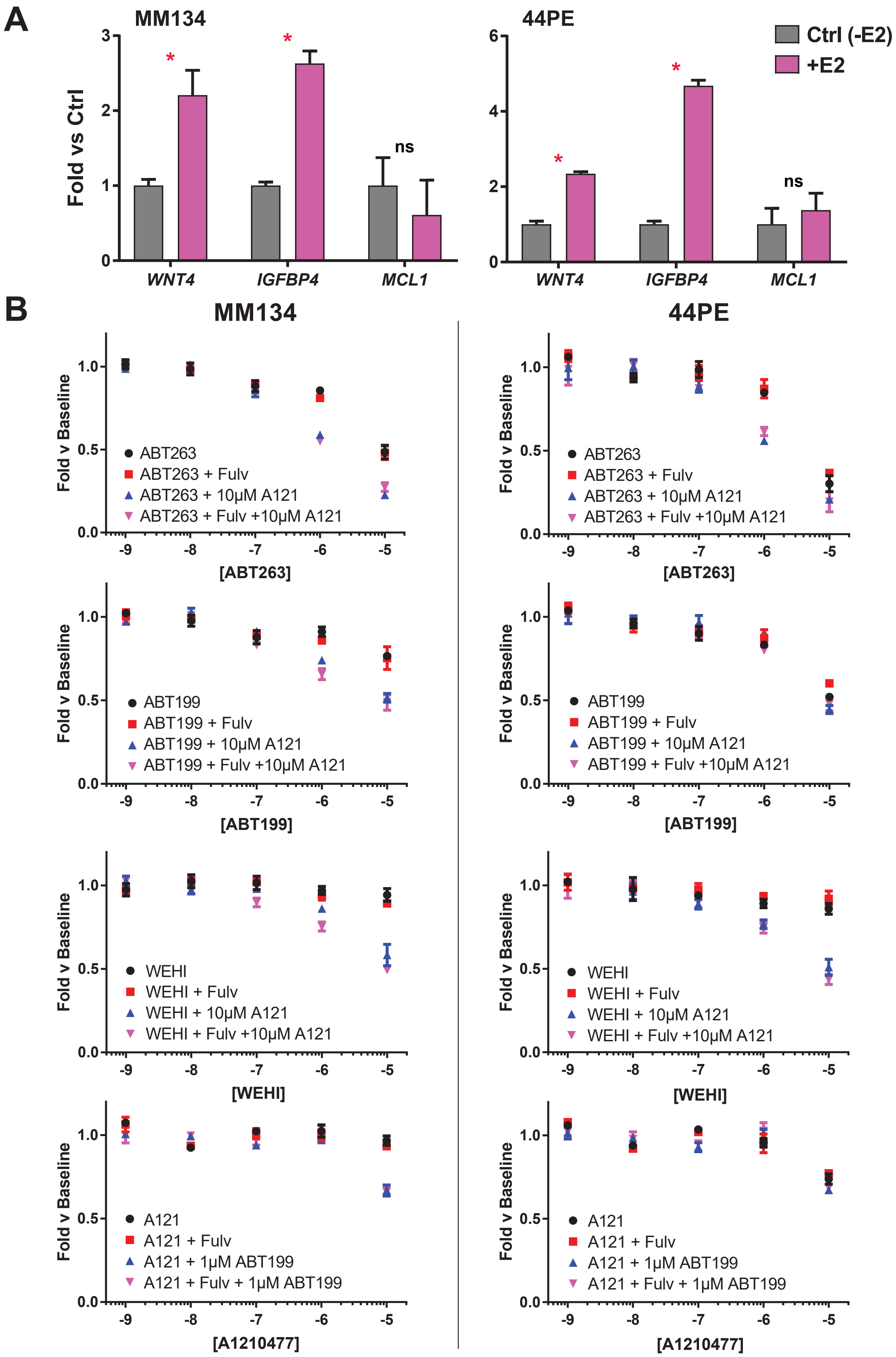
A
MM134
44PE
Fold vs Ctrl
Ctrl (-E2)
+E2
*
ns
WNT4
IGFBP4
MCL1
B
MM134
44PE
Fold v Baseline
ABT263
ABT263 + Fulv
ABT263 + 10μM A121
ABT263 + Fulv +10μM A121
[ABT263]
ABT199
ABT199 + Fulv
ABT199 + 10μM A121
ABT199 + Fulv +10μM A121
[ABT199]
WEHI
WEHI + Fulv
WEHI + 10μM A121
WEHI + Fulv +10μM A121
[WEHI]
A121
A121 + Fulv
A121 + 1μM ABT199
A121 + Fulv + 1μM ABT199
[A1210477]

**Supplemental Figure 4. ER regulation of MCL-1 is not linked to shifts in sensitivity to BH3 mimetic drugs.** (A), Cells were hormone-deprived prior to siRNA reverse transfection and subsequent treatment with 100pM E2 for 24hrs. Expression was normalized to *RPLP0*. *, Ctrl vs +E2, $p < 0.05$, Student's T-test. (B), Cells in full serum were pre-treated with 100nM Fulvestrant and/or base BH3 mimetic (ie. the compound at static concentration per assay) for 6hrs prior to treatment with increasing concentrations of the test BH3 mimetic. Cell proliferation/death was assessed by dsDNA quantification 72hrs after second treatment. Points represent mean of biological triplicate ±SD.

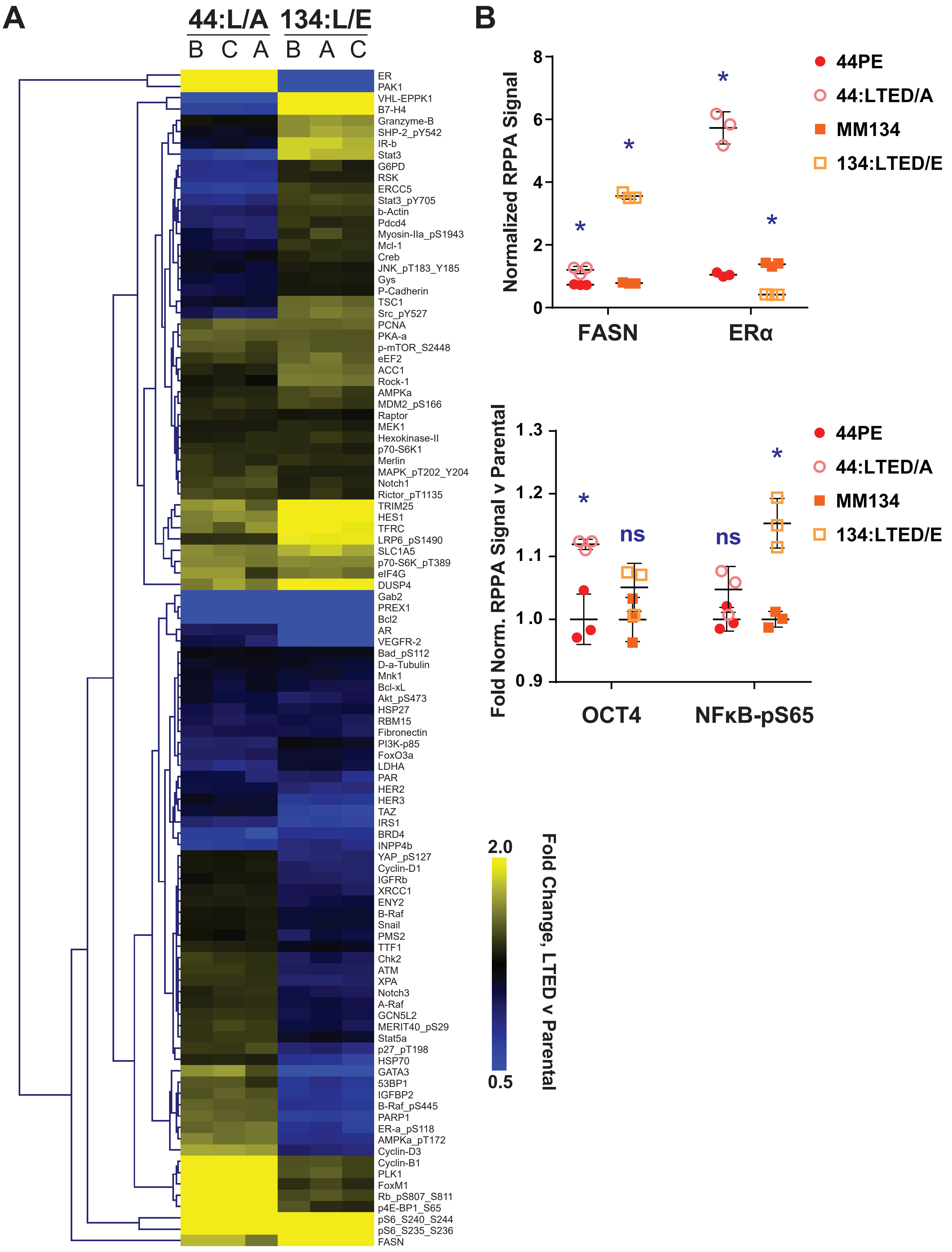

A
44:L/A
134:L/E
B C A
B A C
ER
PAK1
VHL-EPPK1
B7-H4
Granzyme-B
SHP-2_pY542
IR-b
Stat3
G6PD
RSK
ERCC5
Stat3_pY705
b-Actin
Pdcd4
Myosin-IIa_pS1943
Mcl-1
Creb
JNK_pT183_Y185
Gys
P-Cadherin
TSC1
Src_pY527
PCNA
PKA-a
p-mTOR_S2448
eEF2
ACC1
Rock-1
AMPKa
MDM2_pS166
Raptor
MEK1
Hexokinase-II
p70-S6K1
Merlin
MAPK_pT202_Y204
Notch1
Rictor_pT1135
TRIM25
HES1
TFRC
LRP6_pS1490
SLC1A5
p70-S6K_pT389
eIF4G
DUSP4
Gab2
PREX1
Bcl2
AR
VEGFR-2
Bad_pS112
D-a-Tubulin
Mnk1
Bcl-xL
Akt_pS473
HSP27
RBM15
Fibronectin
PI3K-p85
FoxO3a
LDHA
PAR
HER2
HER3
TAZ
IRS1
BRD4
INPP4b
YAP_pS127
Cyclin-D1
IGFRb
XRCC1
ENY2
B-Raf
Snail
PMS2
TTF1
Chk2
ATM
XPA
Notch3
A-Raf
GCN5L2
MERIT40_pS29
Stat5a
p27_pT198
HSP70
GATA3
53BP1
IGFBP2
B-Raf_pS445
PARP1
ER-a_pS118
AMPKa_pT172
Cyclin-D3
Cyclin-B1
PLK1
FoxM1
Rb_pS807_S811
p4E-BP1_S65
pS6_S240_S244
pS6_S235_S236
FASN
2.0
0.5
Fold Change, LTED v Parental
B
Normalized RPPA Signal
8
6
4
2
0
FASN
ERα
*
44PE
44:LTED/A
MM134
134:LTED/E
Fold Norm. RPPA Signal v Parental
1.3
1.2
1.1
1.0
0.9
OCT4
NFκB-pS65
ns

**Supplemental Figure 5. RPPA identifies diverse signaling pathways activated during anti-estrogen resistance in ILC models.** (A), Heatmap for n=104 shared protein signaling changes for LTED versus parental cells. A/B/C represent biological replicate samples. (B), Protein changes in LTED models that support previously described features of the anti-estrogen resistance phenotype in these models. Points represent individual biological replicates. *, Matched parental v LTED, q<0.05.

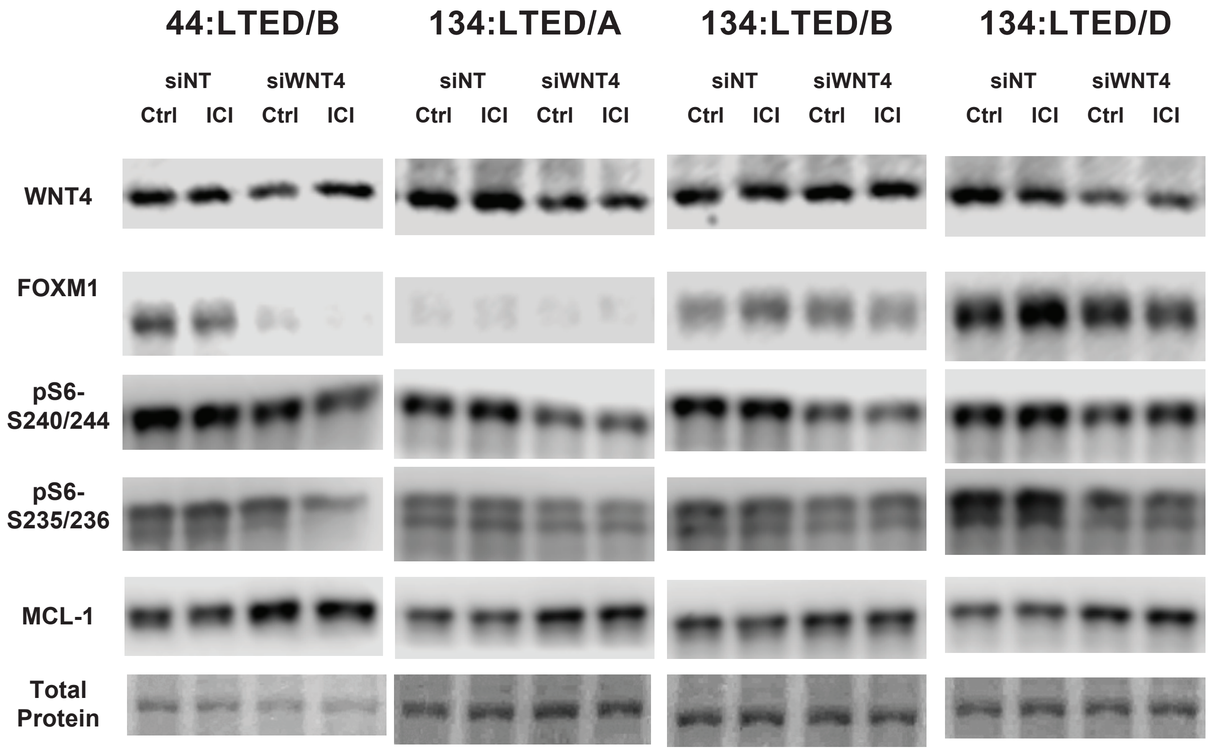


**Supplemental Figure 6. WNT4 regulation of key targets is consistent across ILC LTED cell lines.** LTED cells (in hormone-deprivation) were reverse transfected with the indicated siRNA. 24hrs later, cells were treated with 1μM Fulvestrant or vehicle, and lysates were harvested after an additional 24hrs. Total protein detected with Ponceau staining.

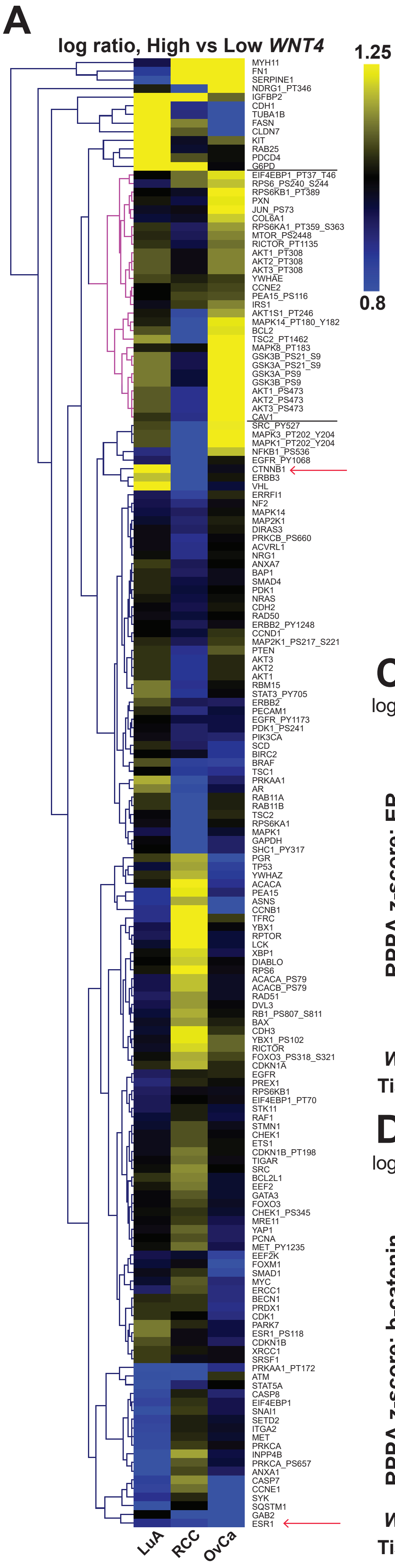


**B** Ovarian (Serous) RPPA, High vs Low *WNT4* mRNA

| Gene | Log Ratio | p-Value | q-Value | Gene | Log Ratio | p-Value | q-Value |
|---|---|---|---|---|---|---|---|
| **GSK3A_PS9** | 0.45 | 3.09E-08 | **3.20E-06** | MTOR_PS2448 | 0.13 | 0.0195 | 0.144 |
| **GSK3B_PS9** | 0.45 | 3.09E-08 | **3.20E-06** | NDRG1_PT346 | 0.39 | 0.0208 | 0.147 |
| **CASP7** | -0.42 | 3.00E-06 | **2.07E-04** | AKT1S1_PT246 | 0.16 | 0.0215 | 0.147 |
| **TSC2_PT1462** | 0.29 | 1.31E-05 | **4.71E-04** | PIK3CA | -0.12 | 0.0221 | 0.147 |
| **GSK3A_PS21_S9** | 0.33 | 1.37E-05 | **4.71E-04** | EGFR_PY1173 | -0.07 | 0.0238 | 0.147 |
| **GSK3B_PS21_S9** | 0.33 | 1.37E-05 | **4.71E-04** | PRDX1 | -0.15 | 0.0243 | 0.147 |
| **SMAD1** | -0.22 | 5.76E-05 | **1.70E-03** | PRKAA1 | -0.11 | 0.0248 | 0.147 |
| **CCNE1** | -0.43 | 1.82E-04 | **4.71E-03** | RPS6KB1_PT389 | 0.25 | 0.0249 | 0.147 |
| **SYK** | -0.49 | 5.31E-04 | **0.0106** | MAPK14_PT180_Y182 | 0.23 | 0.0268 | 0.154 |
| **MAPK8_PT183** | 0.36 | 6.40E-04 | **0.0106** | FOXM1 | -0.25 | 0.036 | 0.197 |
| **AKT1_PS473** | 0.46 | 6.65E-04 | **0.0106** | CDK1 | -0.14 | 0.0369 | 0.197 |
| **AKT2_PS473** | 0.46 | 6.65E-04 | **0.0106** | YWHAZ | -0.18 | 0.0379 | 0.197 |
| **AKT3_PS473** | 0.46 | 6.65E-04 | **0.0106** | ESR1 | -0.54 | 0.0388 | 0.197 |
| **SQSTM1** | -0.55 | 8.18E-04 | **0.0121** | RICTOR_PT1135 | 0.11 | 0.039 | 0.197 |
| **GAB2** | -0.4 | 1.86E-03 | **0.0256** | BRAF | -0.2 | 0.0431 | 0.213 |
| **RPS6KA1_PT359_S363** | 0.15 | 2.10E-03 | **0.0271** | CAV1 | 0.46 | 0.0488 | 0.235 |
| **EEF2K** | -0.2 | 3.05E-03 | **0.0368** | PCNA | -0.1 | 0.0562 | 0.264 |
| **JUN_PS73** | 0.25 | 3.20E-03 | **0.0368** | IRS1 | 0.13 | 0.0574 | 0.264 |
| **CLDN7** | -0.62 | 4.22E-03 | **0.0459** | CCNB1 | -0.35 | 0.0597 | 0.265 |
| **EIF4EBP1_PT37_T46** | 0.22 | 4.72E-03 | **0.0468** | CDH1 | -0.44 | 0.061 | 0.265 |
| **BIRC2** | -0.17 | 4.75E-03 | **0.0468** | BECN1 | -0.14 | 0.0615 | 0.265 |
| **ERCC1** | -0.14 | 5.12E-03 | **0.0482** | COL6A1 | 0.2 | 0.0747 | 0.316 |
| MYH11 | 1.19 | 6.65E-03 | 0.0599 | FASN | -0.27 | 0.0804 | 0.333 |
| CASP8 | -0.12 | 8.30E-03 | 0.0716 | MYC | -0.13 | 0.0828 | 0.336 |
| PXN | 0.23 | 0.0108 | 0.0892 | PEA15 | -0.12 | 0.0918 | 0.359 |
| BCL2 | 0.22 | 0.0144 | 0.111 | SRC_PY527 | 0.24 | 0.092 | 0.359 |
| ASNS | -0.26 | 0.0145 | 0.111 | RPS6_PS240_S244 | 0.19 | 0.0943 | 0.362 |

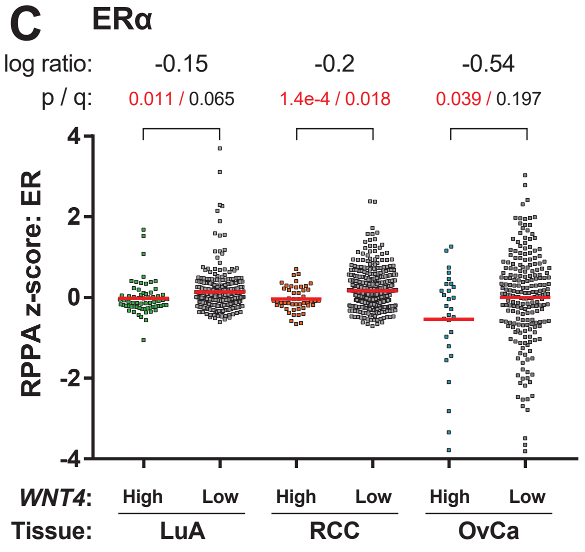


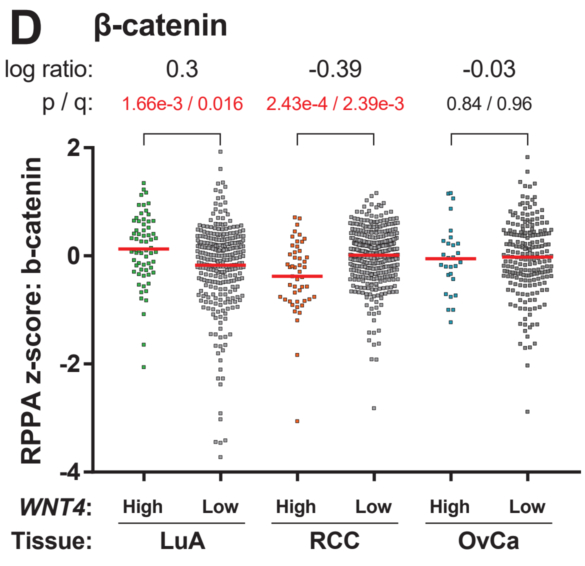


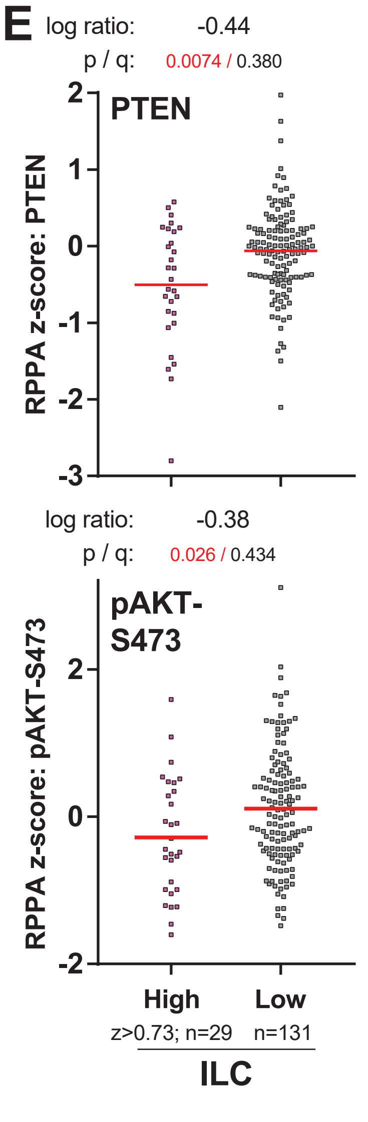

**F**

| **Signature** | **pvalue** | **qvalue** |
|---|---|---|
| HALLMARK_PROTEIN_SECRETION | 6.52E-05 | 0.009953 |
| CTIP_DN.V1_DN | 8.37E-05 | 0.009953 |
| HALLMARK_MTORC1_SIGNALING | 0.000135 | 0.010696 |
| TBK1.DF_DN | 0.000372 | 0.022107 |

**Supplemental Figure 7. High *WNT4* expression is associated with distinct signaling pathways in varying tumor types.** (A), TCGA RPPA data from Figure 5B; n=170 protein targets different in high v low *WNT4*-expressing tumors in at least one tumor type at p<0.1. Brackets identify mTOR signaling targets increased only in OvCa; arrows denote targets in (C-D). (B), All protein target changes in OvCa in high v low *WNT4*-expressing tumors at p<0.1; bold denotes q<0.05. (C-D), RPPA z-score for shared signaling components discussed in text. Colored v Gray points represent individual tumor samples with high/low *WNT4* from (A). Line represents mean RPPA signal for the indicate group. Log ratio shown as High v Low *WNT4*. (E), No protein signals at q<0.05 were identified in *WNT4* high v low expressing ILC; PTEN and pAkt-S473 were included among n=17 differentially expressed protein targets at p<0.05. Colored v Gray points represent individual tumor samples with high/low *WNT4*. (F), TCGA RNAseq data from ER+ breast cancers (n=814) were used to identify genes with expression correlated with  WNT4. The top 500 genes (q<5e-9) were used for over-representation analysis against the MSigDB Hallmark, C2/Canonical Pathways, and C6 datasets.

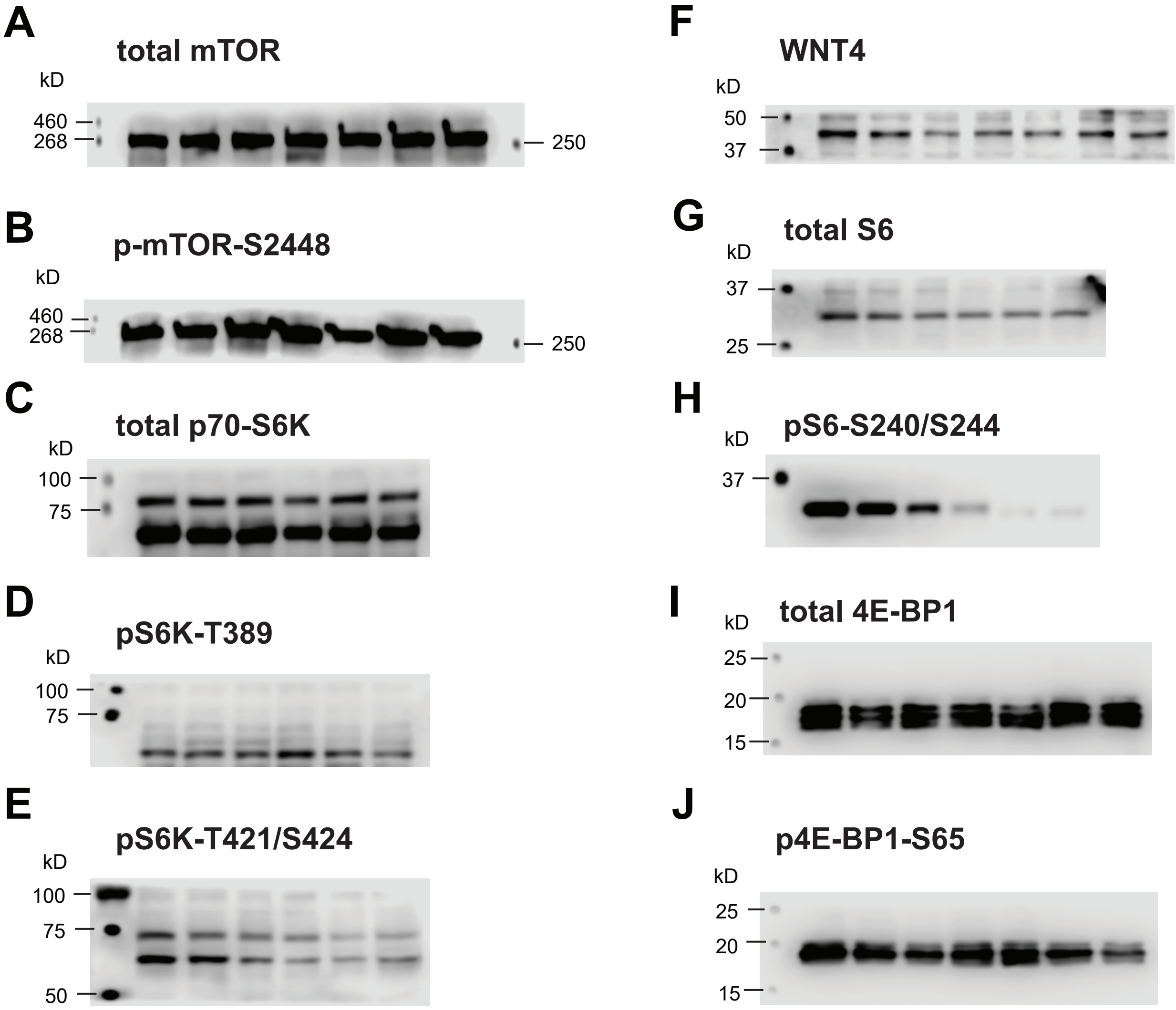


**Supplemental Figure 8. Molecular weight markers for immunoblots.**
Immunoblot images above are derived from figures herein (as noted for each sub-panel) and are representative of all related images. Dots that are labeled as weight markers are from marking the molecular weight ladder with a WesternSure pen prior to imaging. Antibodies are listed in Materials and Methods.
(A), Figure 3A - mTOR; predicted weight ~289kD. (B), Figure 3A - p-mTOR-S2488; predicted weight ~ 289kD. (C), Figure 3B - total p70-S6K; predicted weight ~ 70, 85kD. (D), Figure 3C - pS6K-T389; predicted weight ~ 70, 85kD. (E), Figure 3C - pS6K-T421/S424; predicted weight ~ 70, 85kD. (F), Figure 3A - WNT4; predicted weight ~ 39kD. (G), Figure 3B - total S6; predicted weight ~32kD. (H), Figure 3B - pS6-S240/S244; predicted weight ~ 32kD. pS6-S235/S236 also has a predicted weight of ~ 32kD. (I), Figure 3A - total 4E-BP1, predicted weight ~ 15 - 20kD. (J), Figure 3A - p4E-BP1-S65; predicted weight ~ 15 - 20kD.
Exposure was increased (vs Figures 3A and 3B) to show ladder makings for (A), (B), (C), (H), (I), and (J). Exposure was decreased (vs Figure 3C) to show ladder markings for (E).
