## Supplemental Document 1 for "Estrogen controls mTOR signaling and mitochondrial function via WNT4 in lobular carcinoma cells"

48 samples probed with
305 antibodies

RPPA Set123

Aug 31 2016

Your samples are arranged in RPPA CORE 08082016_123 (Set123).
If you have any questions regarding your samples or data, please refer to the set number.

Processed by: Ling, Doris

### PROCEDURES FOR RPPA CORE 08082016_123 (Set123).

1) Tumor or cell lysates were serially diluted two-fold for 5 dilutions (from undiluted to 1:16 dilution) and arrayed on nitrocellulose-coated slides in an 11x11 format.

2) Samples were probed with antibodies by tyramide-based signal amplification approach and visualized by DAB colorimetric reaction.

3) Slides were scanned on a flatbed scanner to produce 16-bit tiff image.

4) Spots from tiff images were identified and the density was quantified by Array-Pro Analyzer.

5) Relative protein levels for each sample were determined by interpolation of each dilution curves from the "standard curve" (supercurve) of the slide (antibody). Supercurve is constructed by a script in R, written by Bioinformatics. These values (given as Log2 values) are defined as Supercurve Log2 (Raw) values and shown in the Excel worksheet tab labeled "RawLog2".

6) All the data points were normalized for protein loading and transformed to linear value, designated as "Normalized Linear" (labeled "NormLinear" in the worksheet).
*The linear value can be used for bar graphs or further analyses according to your study design.
We have recently implemented an improved normalization algorithm for protein loading correction and antibody variation adjustment. This approach is critical to provide accurate values for RPPA data merging if you have samples performed in separate RPPA sets that you wish to combine. We listed the protein loading correction factors (CF1 and CF2) in the last column on this page (NormLinear) for your reference. Each sample has its unique correction factor. CF1 values are calculated among your sample set while CF2 values are calculated among the entire 1056 samples on the same slide. In contrast to CF1 indicating protein-loading factor among your sample set, CF2 determines protein level for each sample in the entire set of RPPA. If the correction factor is less than 0.25 or greater than 2.5, we consider these samples “outliers,” indicating that protein concentration is much lower or much higher than the other samples. We suggest that you exclude these “outliers” from further analysis. (One exception: intrinsic protein expression patterns can skew

the correction factor to some extent. In that case, you may want to include this data, but we suggest that you examine the data set carefully.)

7) "Normalized Linear" values were transformed to Log2 values (labeled "NormLog2" in worksheet), and then median-centered for hierarchical clustering analysis (labeled "NormLog2_MedianCentered" in the worksheet). Median-centered values were then formatted for heatmap generation in the "Format for Heatmap" worksheet.

8) We included heatmaps for an unsupervised hierarchical cluster (unsupervised on both antibodies and samples) and an antibody unsupervised but samples are arranged in the order you submitted them, for your reference.

9) The heatmap included was generated in Cluster 3.0 (http://bonsai.hgc.jp/~mdehoon/software/cluster/software.htm) as a hierarchical cluster using Pearson Correlation and a center metric. The resulting heatmap was visualized in Treeview {http://www.eisenlab.org/eisen/) and presented as a high resolution .bmp format.

10) We stained RPPA slides for 305 unique antibodies, which were analyzed, on Array-Pro then by SuperCurve Rx64 3.1.1. We performed QC tests for each antibody staining (slide). The QC Score (Probability) values were also included in the data spreadsheet for your reference. A QC score above 0.8 indicates good antibody staining. We only included the data for the 305 individual antibodies with QC Scores higher than 0.80 in the heatmaps.

For samples of tissue origin, we removed 5 antibodies that cross-reacted with "damaged components" (of unknown mechanism). There are 61 individual mouse and rat antibodies (labeled "-M" and "-T") that will be removed from mouse tissue, rat tissue, or xenograft samples, unless you specifically indicated for inclusion in your Sample Submission Form.

11) The bioinformatics should be done at your end from the Excel file we provide. We recommend that you create bar graphs based on the data in the Excel file. The heatmaps are solely to provide overall patterns.

Antibody status for RPPA
(V) = Validated antibody for RPPA
(C) = Use with caution. Validation in progress

(Q) = These antibodies recognize unidentified "damaged" component(s) in addition to its specific protein. The "damaged" component(s) were observed only in certain tissue samples.
(E) = Under Evaluation
(M) = Mouse antibody was used
(G) = Goat antibody was used
(R) = Rabbit antibody was used
(T) = Rat antibody was used

**Example: Akt_pS473-R-V_GBL9016996 means the antibody specifically recognizes Akt phosphorylated on Serine 473. This is a rabbit antibody validated for RPPA application. GBL9016996 is the slide ID (barcode). The slide ID is not included in the final version of heatmap.

<u>**Your samples are arranged in RPPA CORE 08082016_123 (Set123). If you have any questions regarding your samples or data, please refer to the set number in your inquiries.**</u>

### Important Information for RPPA Report

Thank you for using the RPPA Core for your functional proteomics studies. Your RPPA results are provided in an Excel file with multiple pages (tabs).

A. The 1st page (RawLog2) provides the raw RPPA data in Log2 values without any normalization.

B. The 2nd page (NormLinear) provides normalized RPPA data in linear values, which can be used for bar graphs or line graphs. **We have recently implemented an improved normalization algorithm for protein loading correction and antibody variation adjustment.** This approach is critical to provide accurate values for RPPA data merging if you have samples performed in separate RPPA sets that you wish to combine. We listed the protein loading correction factor (CF) in the last column on this page (NormLinear) for your reference. Each sample has its unique correction factor. If the correction factor is less than 0.25 or greater than 2.5, we consider these samples “outliers,” indicating that protein concentration is much lower or much higher than the other samples. We suggest that you exclude these “outliers” from further analysis. (One exception: intrinsic protein expression patterns can skew the correction factor to some extent. In that case, you may want to include this data, but we suggest that you examine the data set carefully.)

C. The 3rd page (NormLog2) provides normalized RPPA data in Log2 values for further bioinformatics analysis.

D. In order to make heatmaps for data visualization, we processed for median centering across each antibody from normalized Log2 values. These values are presented on the 4th page (NormLog2 Median Centered).

E. Beware of outliers in the heatmaps. They can be determined by viewing the values in the included Excel spreadsheet under tab name “NormLinear" and the last column (CF). If the values in this column are < 0.25 or > 2.5, they are considered outliers.

We provide two heatmaps for your reference: (1) unsupervised hierarchical clustering and (2) supervised by the sample order according to your design. “Red” in the heatmaps means above median and “green” means below median.

Please note that your RPPA data report is presented in Excel format. Heatmaps display visualization figures for your reference. Further bioinformatics analysis should be performed at your end.

The heatmaps were developed by the MD Anderson Cancer Center Department of Bioinformatics and Computational Biology, In Silico Solutions, Santeon and SRA International. This work was supported in part by U.S. National Cancer Institute (NCI; MD Anderson TCGA Genome Data

Analysis Center) grant numbers CA143883 and CA083639, the Mary K. Chapman Foundation, the Michael & Susan Dell Foundation (honoring Lorraine Dell), and MD Anderson Cancer Center Support Grant P30 CA016672 (the Bioinformatics Shared Resource).

Functional Proteomics RPPA Core facility is supported by MD Anderson Cancer Center Support Grant # 5 P30 CA016672-40. Publications using data generated by the Core should cite the Core grant in the acknowledgement section.

### Important Information for RPPA Antibody List

Reverse phase protein arrays (RPPA) are dependent on the quality of antibodies and control metrics used for each antibody and each sample. We perform extensive evaluation of each antibody and also have a continuous reassessment program to ensure delivery of high quality data.

It is important to emphasize that although the majority of our antibodies are from commercial sources and usually are monoclonal antibodies, behavior of the antibodies can change over time and also between batches. Furthermore, with ongoing use and reassessment on our RPPA platform, new information comes to light (particularly under different conditions and with different types of samples) that may result in a change in validation status and raise concerns about the utility of specific antibodies under particular conditions. We are committed to providing high quality data and thus will update the description of the antibody performance on our RPPA website. http://www.mdanderson.org/education-and-research/resources-for-professionals/scientific-resources/core-facilities-and-services/functional-proteomics-rppa-core/index.html.

We will provide updates on major concerns by e-mail; however, we recommend that you frequently visit our web page to review the antibody list and validation status of each antibody. The antibodies are labeled as follows: "valid" which means they perform well in all assays available; "use with caution" means that under most circumstances the antibodies provide high quality information but we have concerns that they may not perform well under some conditions; and "under evaluation" means that the antibodies are currently being evaluated or re-evaluated for performance. We would like to emphasize that RPPA is best thought of as a high throughput screening assay. We recommend that all RPPA results be confirmed by an orthologous approach.

The antibodies listed on our website perform well both in cell lines and patient samples based on the designations listed above. Any antibodies that are not on our current list may have challenges. If you have antibodies that are no longer listed, we recommend that you consider the data with caution, and contact the RPPA Core to determine the reason for removal from our standard list. You may also contact us on the status of any specific antibodies of interest not listed.

### Antibodies that do not pass current quality control should be deleted from previous data sets

We have identified a set of antibodies where their performance no longer meets our standards as they have been shown to have liabilities under a number of conditions.

These antibodies include:

- AIB1 (BD Biosciences #611105)
- Caspase-9_cleavedD330 (Cell Signaling Technology #9501)
- COX2 (Epitomics #2169-1)
- IGF-1R-beta (Cell Signaling Technology #3027)
- PTCH (SDI 2113.0002)
- TAZ_pS89 (Santa Cruz #sc-17610)
- VASP (Cell Signaling Technology #3112)

We no longer run these antibodies in RPPA, and will continue searching for alternative sources. If you have these antibodies in your previous data set, we recommend that you remove them from further consideration.

### Antibodies that cross-react within EGFR family members

As a further part of our analysis, we have found that a number of antibodies to phospho-HER2 and phospho-EGFR cross-react when the opposite molecule is present at very high levels. However, they perform well when the other receptor is not highly expressed so we will continue to provide the information.

**These antibodies include:**

A. EGFR antibody from Santa Cruz (#sc-03) cross-reacts significantly with overexpressed HER2. (We have removed this antibody from our list.)
B. Phospho-EGFR Y1068 (Cell Signaling Technology, #2234), and phospho-EGFR Y992 antibodies (Cell Signaling Technology, #2235) modestly cross-react with overexpressed phosphorylated HER2.
C. Similarly, the phospho-HER2 Y1248 antibody from Millipore (#06-229) modestly cross-reacts with phospho-EGFR.

In contrast, we have demonstrated a set of non-cross-reacting antibodies against EGFR family members under our test conditions. We have included these antibodies in our recently revised antibody list. These antibodies include:

A. EGFR antibody from Cell Signaling Technology (#2232) does not demonstrably cross-react with HER2.
B. HER2 antibodies from Lab Vision (#MS-325-P1) and Epitomics (#2064) do not demonstrably cross-react with EGFR.
C. The phospho-EGFR Y1173 antibody from Epitomics does not detectably cross-react with any other EGFR family member including HER2.

### Antibodies cross react with “damaged components” in tissue samples

As part of our continuous reassessment process, we have identified a series of antibodies that may not perform well in patient samples. We are concerned that these antibodies may cross-react with “damaged components” present in patient samples. While we do not understand the mechanism leading to increased cross-reacting levels of such components in a subset of patient samples, we recommend that **tissue samples** containing coordinately high levels of these components identified by such antibodies be removed from further consideration. We recommend that these antibodies not be used further in analysis of patient samples other than as quality control.

**These antibodies include:**

- c-Met (Cell Signaling Technology #3127)
- ERCC1 (Lab Vision #MS-671-PO) (Not performed in current RPPA analysis)
- Caspase 8 (Cell Signaling Technology #9746)
- Cleaved PARP (Cell Signaling Technology #9564)
- Rab25 (Covance Custom) (Not performed in current RPPA analysis)
- Rb (Cell Signaling Technology #9309)
- SETD2 (Abcam #ab69836)
- Smac (Cell Signaling Technology #2954)
- Snail (Cell Signaling Technology #3895)

**Please note that we designate these antibodies as “Used for QC” in our antibody list. We do report these antibodies in cell line samples but not in tissue samples.

### Protein phosphorylation in tissue samples may change during sample processing

We have evaluated and reported that phosphorylation of certain specific proteins (especially the EGFR/MAPK signaling module) in tissue can be altered by cold ischemia during sample processing, while levels of total protein and the majority of phosphoproteins remain constant. Although the global proteome is remarkably stable, we still recommend that every effort be made to limit the time from tissue excision to sample preservation, aiming to reduce modifiable pre-analytical variables.

<u>Please refer to the following references regarding phosphoproteins susceptible to tissue sample processing.</u>

A) Hennessy, B. and Lu, Y. et al, A technical assessment of the utility of reverse phase protein arrays for the study of the functional proteome in non-microdissected human breast cancers. Clinical Proteom 6:129-151, 2010. PMID 21691416.

B) Mertins, P. and Yang, F. et al, Ischemia in tumors induces early and sustained phosphorylation changes in stress kinase pathways but does not affect global protein levels. Molecular and Cellular Proteomics, May 7, 2014. In Press. (Manuscript M113.036392, PMID: 24719451)

### In Summary

We provide RPPA data of your 48 samples probed with 305 antibodies. The data set is arranged in Excel format, while two heatmaps are also provided for your visualization of each dataset.

Please review your data set with 48 samples carefully, especially on the 2nd page (NormLinear) of Excel file. We highlighted those samples having problems with “protein loading” and considered “outliers” on your study or their protein expression patterns skewing the correction factors. Please interpret this data with caution.

### Supervised clustering

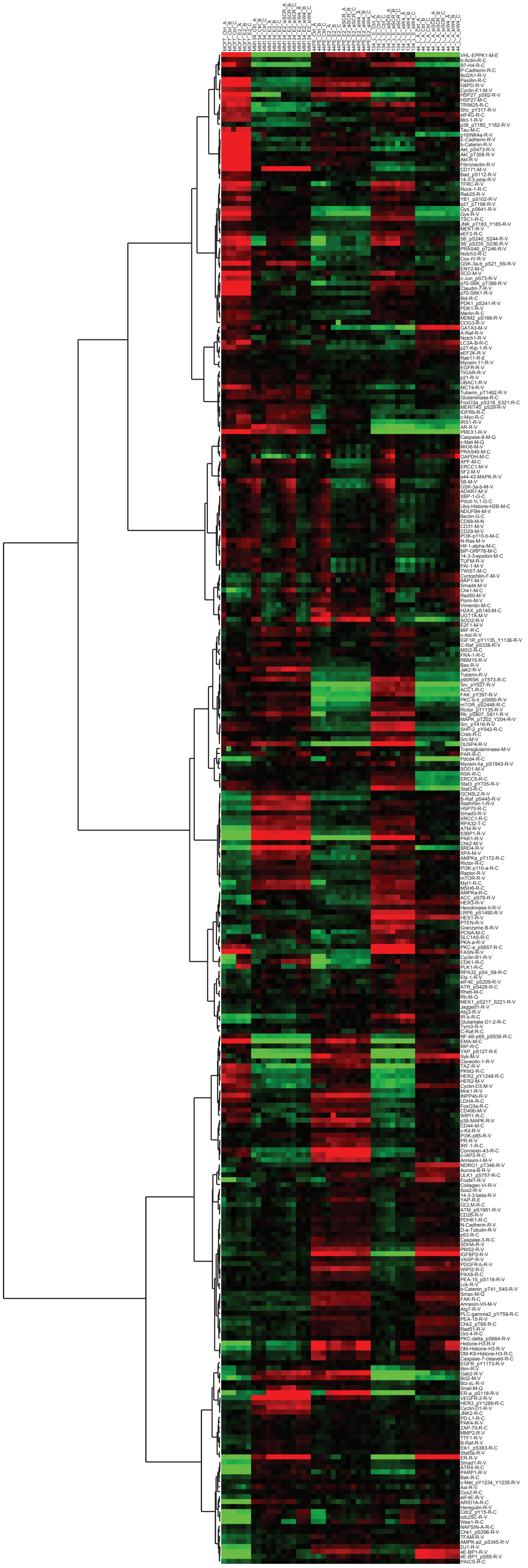

Unsupervised clustering

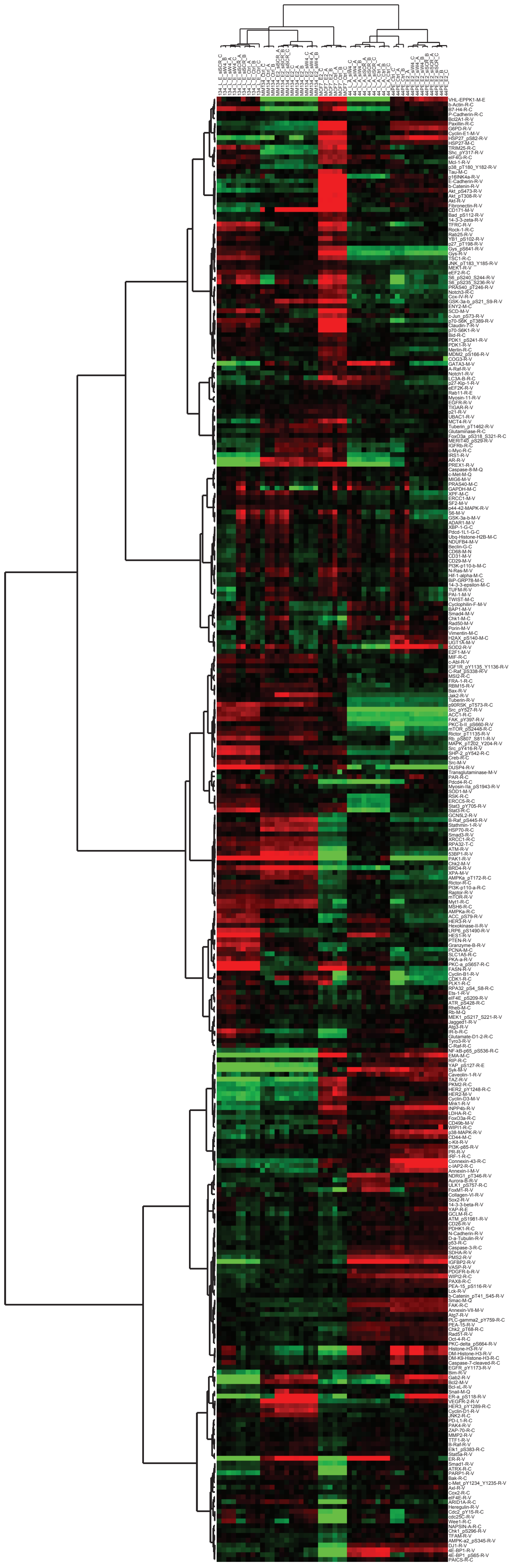
